# An age-specific platelet differentiation path from hematopoietic stem cells contributes to exacerbated thrombosis

**DOI:** 10.1101/2024.02.23.581812

**Authors:** DM Poscablo, AK Worthington, S Smith-Berdan, BA Manso, R Adili, T Cool, RE Reggiardo, S Dahmen, AE Beaudin, SW Boyer, M Holinstat, EC Forsberg

**Affiliations:** Institute for the Biology of Stem Cells, University of California-Santa Cruz, Santa Cruz, CA, USA; Program in Biomedical Sciences and Engineering, Department of Molecular, Cell, and Developmental Biology, University of California-Santa Cruz, Santa Cruz, CA 95064, USA; Current affiliation: Departments of Internal Medicine and Pathology, and Program in Molecular Medicine, School of Medicine, University of Utah, Salt Lake City, UT 84112, USA; Department of Pharmacology, University of Michigan Medical Center, Ann Arbor, MI 48109, USA; Biomolecular Engineering, University of California-Santa Cruz, Santa Cruz, CA, USA

## Abstract

Platelet dysregulation is drastically increased with advanced age and contributes to making cardiovascular disorders the leading cause of death of elderly humans. Hematopoietic stem and progenitor cells continuously give rise to platelets, but their contributions to variable platelet production and activity throughout life remain unclear. Here we reveal a direct differentiation pathway from hematopoietic stem cells into platelets that is unique to aging. An unequivocal genetic lineage tracing mouse model demonstrated that this age-specific pathway is progressively propagated over time. Remarkably, the age-specific platelet path is decoupled from all other hematopoietic lineages, including erythropoiesis, and operates as an additional layer in parallel with canonical platelet production. This results in two molecularly and functionally distinct populations of megakaryocyte progenitor cells that that operate in parallel. The age-specific megakaryocyte progenitor population has profoundly enhanced capacity to engraft, expand, and reconstitute platelets, and produces an additional platelet population that exists only in old mice. Consistent with increased thrombotic incidence upon aging, the two pools of co-existing platelets contribute to age-related thrombocytosis and dramatically increased thrombosis *in vivo*. Upon acute, platelet-specific stress, the age-specific MkPs endowed old mice with superior capacity to rapidly restore platelet counts. These findings reveal stem cell-based aging as a mechanism for platelet dysregulation and identify an aging-induced population of functionally enhanced MkPs as a unique source of age-specific platelets.

**>HIGHLIGHTS:**

- Aging leads to two parallel platelet specification paths from HSCs
- The shortcut platelet pathway is perpetuated by highly expansive MkPs unique to aging
- The age-specific differentiation path contributes to thrombosis and platelet hyperreactivity
- Age-specific MkPs serve as potent first responders to acute platelet loss

## INTRODUCTION

Aging is characterized by the stereotypical decline in tissue function and is the primary risk factor for major diseases. In particular, while platelets (Plts) are critical for controlled hemostasis, their dysregulated production and activation during aging contribute to the pathogenesis of platelet-related disorders in the elderly. Tipping the homeostatic balance towards inadequate Plt production is associated with higher risk for bleeding disorders, whereas Plt overproduction and hyperactivation lead to pathologic clot formation in thrombotic diseases such as deep vein thrombosis and ischemic stroke (Kim et al. 2006; Dayal et al. 2013; Gleerup & Winther 1995; Virani et al. 2020; Moulis et al. 2014; Neylon et al. 2003; Yang et al. 2016; Davizon-Castillo et al. 2019). Plts are extremely short-lived (∼4 days in mice and ∼8 days in humans) and therefore continuously derived from hematopoietic stem cells (HSCs) in the bone marrow (BM) via a gradual differentiation cascade of intermediate progenitor cells towards megakaryocyte progenitors (MkPs) (Noetzli et al. 2019). While the differentiation path from HSCs to MkPs may be altered upon in vitro or in vivo stress and upon aging (Roch et al. 2015; Sanjuan-Pla et al. 2013a; Haas et al. 2015), we and others have shown that in young, unperturbed mice, MkPs and Plts, like all other adult hematopoietic lineages, arise via a Flk2+ differentiation stage (**Figure 1A**, t**op row of Figure 1B**) (Boyer et al. 2011; Buza-Vidas et al. 2011). HSCs in the young adult “FlkSwitch” mouse model express Tomato (Tom), whereas multipotent progenitors (MPPs) and downstream progenitor and mature cells irreversibly switch to GFP expression due to Cre-mediated excision (“floxing”). The FlkSwitch model labels several cell populations with combined platelet/erythroid potential, and we have demonstrated that Plts are GFP+ during development regardless of whether they are derived via Tom+ fetal HSCs or co-existing, developmentally restricted GFP+ HSCs (Beaudin et al. 2016; López et al. 2022; Forsberg et al. 2006; Boyer et al. 2012; Boyer et al. 2011). Analogous differentiation paths and types of identifiable progenitor populations can also be discerned in humans and persist throughout ontogeny (Majeti et al. 2007; Notta et al. 2016). Several independent studies have contributed to the consensus conclusion that hematopoietic aging is characterized by the deterioration of HSC function, most notably demonstrated by their reduced reconstitution capacity compared to young HSCs (yHSCs) (Morrison et al. 1996; Rossi et al. 2008; Sudo et al. 2000; Dykstra et al. 2011). This age-related functional decline of old HSCs (oHSCs) stands in stark contrast to our recently uncovered surprising gain of function of old MkPs (oMkPs) compared to young MkPs (yMkPs) (Poscablo et al. 2021; Poscablo & Forsberg 2021). This led us to hypothesize that age-related alterations in MkPs contribute to dysregulation of Plts in the elderly. Here, we employed the FlkSwitch model as a powerful tool for tracking hematopoietic differentiation pathways to demonstrate a novel and unexpected cellular mechanism for the hematopoietic etiology of Plt-related disorders upon aging.

**Figure 1.**
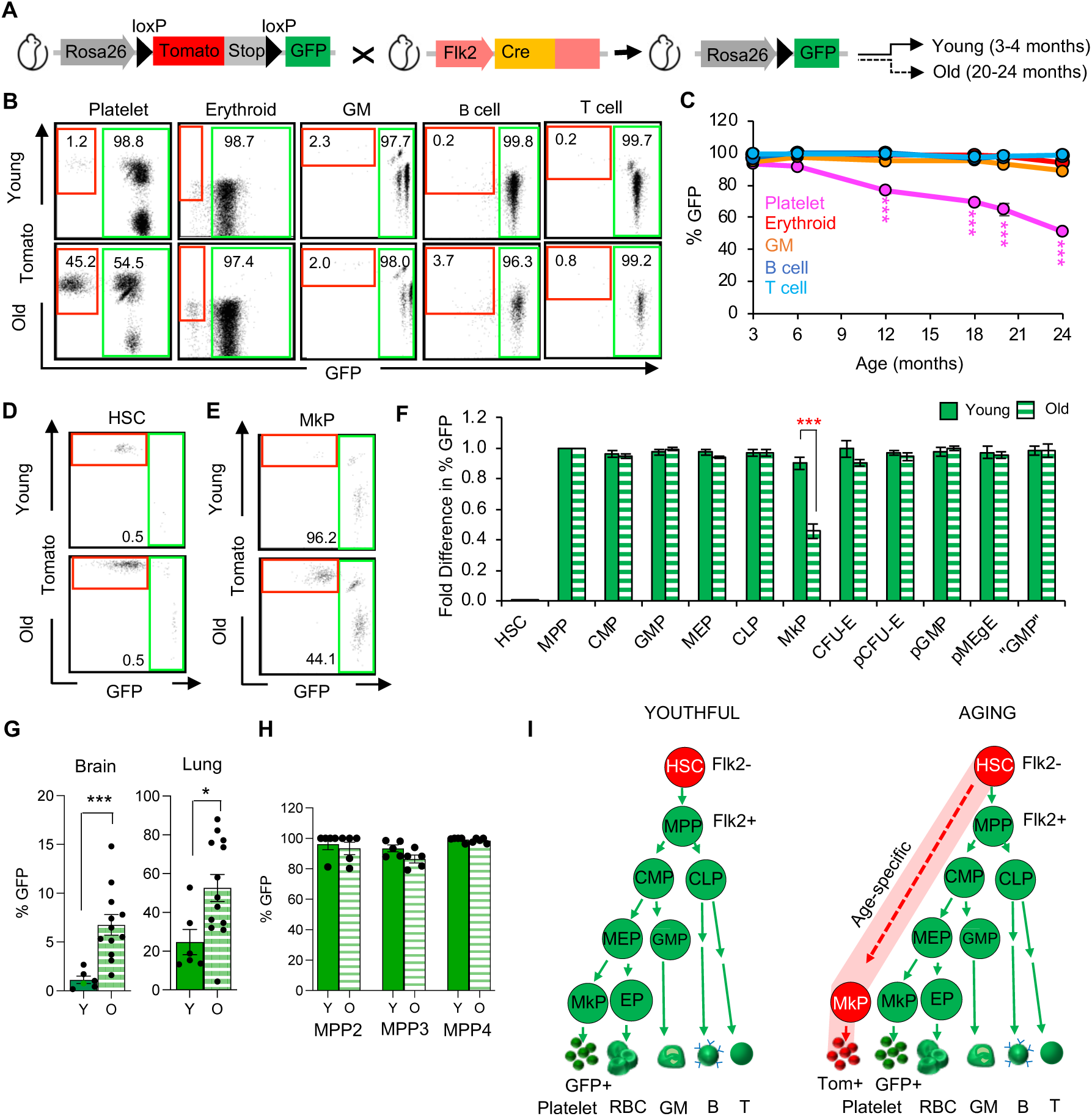
Aging of FlkSwitch mice leads to novel populations of Tomato+ Megakaryocyte Progenitors and Platelets unique to old mice. **A.** Schematic of the mTmG and Flk2-Cre constructs that serve as the basis for the color composition of hematopoietic cells in the FlkSwitch mouse model. Expression of Cre in Flk2+ cells leads to irreversible deletion of Tomato and a switch in expression to GFP in all descendent cells. **B.** Young FlkSwitch mice show very high, equal floxing of mature cells (3 months of age, top row), whereas Tom+ Plts, but not erythroid, GM, B or T cells, increase in aged mice (24-month old, bottom row). **C.** The proportion of GFP+ Plts in the peripheral blood (PB) progressively decrease beyond 12 months of age, whereas the vast majority of erythroid, GM, B and T cells remain GFP+ for life. Quantification of data from B and additional intervening time points are shown. Data represent mean ± SEM of 6 independent experiments, n = 21 mice. Statistics: unpaired t-test compared to baseline 3 months old. ***P<0.0005 **D.** HSCs remain Tom+ for life. Tom versus GFP expression in young and old HSCs. **E.** MkPs are GFP+ in young mice and a novel population of Tom+ MkPs is observed in old mice. Tom versus GFP expression in young and old MkPs. **F.** All progenitors except MkPs maintain efficient switching to GFP-expression throughout life. Fold difference in percent GFP+ classical myeloid, erythromyeloid, and lymphoid progenitor cells compared to MPPs in the BM of young and old FlkSwitch mice. Complete gating strategies are shown in Figure S1. **G.** Tissue resident macrophage GFP-labeling in the FlkSwitch mice increases across tissues with age. Percent GFP labeling of brain microglia and lung alveolar macrophages, defined as in Figure S2. Data represent mean ± SEM of 3 independent experiments, n = 6 young mice, n = 12 old mice. Statistics: t-test. *P<0.05, **P<0.005 **H.** Multipotent progenitor subfractions in the FlkSwitch mice remain GFP+ during aging. Percent GFP labeling of MPP2, MPP3, and MPP4, gated as in Figure S3. Data represent mean ± SEM of 3 independent experiments, n = 5 young mice, n = 5 old mice. Statistics: t-test. Comparisons of Y to O were not statistically different. **I.** Age-specific megakaryopoiesis. Schematic of youthful differentiation pathways in FlkSwitch mice (top) and altered megakaryopoiesis in old FlkSwitch mice (bottom). In young adult and old FlkSwitch mice, HSCs express Tom. Tom is excised upon Cre expression, resulting in GFP+ progenitor and mature cells. Only aged mice have Tom+ MkPs and Plts; cells of all other hematopoietic lineages remain GFP+ throughout life. Tom, Tomato; Y, Young; O, Old; HSC, Hematopoietic Stem Cells; MkP, Megakaryocyte Progenitors; MPP, Multipotent Progenitors; HSPC, Hematopoietic Stem and Progenitor Cells; CMP, Common Myeloid Progenitors; GMP, Granulocyte/Macrophage Progenitor; MEP, Megakaryocyte-erythroid Progenitors; CLP, Common Lymphoid Progenitor; Alternatively defined erythromyeloid progenitors include CFU-E pCFU-E, pGMP, pMEgE, “GMP”; EPs, erythroid progenitor cells; RBCs, red blood cells, GM, granulocytes/macrophages.

## RESULTS

### Platelets and Megakaryocyte Progenitors Uniquely Diverge from a Flk2-Dependent Differentiation Pathway During Aging

To determine the potential changes to lineage specification during aging, we aged FlkSwitch mice and tracked Tom+ and GFP+ hematopoietic cells over time. As previously reported, all mature cell populations in the peripheral blood (PB) of young mice were predominantly GFP+ (Boyer et al. 2011). Surprisingly, aging led to the distinct production of Tom+ Plts, but not erythroid, granulocyte/macrophage (GMs), B or T cells (**Figure 1B**). The proportion of GFP+ Plts in the PB progressively decreased beyond 12 months to reach ∼50% of the total Plt pool by 20+ months in each individual mouse analyzed, whereas the vast majority of erythroid, myeloid (GM), B, and T cells remained GFP+ for life (**Figure 1C**). Examination of bone marrow (BM) populations revealed high fidelity of the FlkSwitch paradigm established with young mice(Boyer et al. 2011; Buza-Vidas et al. 2011): the vast majority of HSCs remained Tom+ for life, and Flk2+ MPPs efficiently switched to GFP expression (**Figure 1D,F**, **Table 1**). Moreover, the overwhelming majority of classical myeloid progenitor cells, including common myeloid progenitors (CMPs), granulocyte/macrophage progenitors (GMPs), megakaryocyte-erythroid progenitors (MEPs), and erythroid progenitors (EPs) were GFP+, as were myeloid populations using alternative markers (**Figure 1F**, **Table 1, Figure S1**) (Pronk et al. 2007). A striking exception was observed for megakaryocyte progenitors (MkPs): nearly half of MkPs in old, but not young, mice remained Tom+ (**Figure 1E-F)**. The divergence of MkP/Plt generation from erythroid production was particularly surprising, as these lineages share critical molecular regulators as well as several progenitor populations with combined erythroid and Plt repopulation capacity (Boyer et al. 2019; Akashi et al. 2000; Yamamoto et al. 2013; Oguro et al. 2013). The uniqueness of this reduced floxing pattern was further underscored by investigation of tissue-resident macrophages (trMacs). Brain and lung trMacs are known to be minimally labeled in young adult FlkSwitch and other Flk2-driven lineage tracing mice (Leung et al. 2019; Bain et al. 2016; Gomez Perdiguero et al. 2014). Interestingly, GFP labeling of trMacs significantly increased in aged FlkSwitch mice (**Figure 1G, Figure S2**), contrasting the Flk2-divergent specification of Plts during aging. The megakaryopoiesis-specific shortcut was observed in every individual mouse that we have aged and analyzed to date (**Table 1**). Analysis of multipotent progenitor pools believed to exist between HSCs and Flk2+ MPPs failed to identify a clear candidate intermediate (**Figure 1H, Figure S3)** (Pietras et al. 2015) indicating that the Tom+ MkPs may derive directly from the Tom+ aged HSCs. Consistent with this idea, only HSC and MkP pools are significantly expanded upon aging (**Figure S1D**) (Valletta et al. 2020; Rundberg Nilsson et al. 2016; Poscablo et al. 2021). Together, these data demonstrate that the canonical Flk2+ differentiation programs of all lineages are robustly maintained throughout life, and that the Plt lineage uniquely deviate from the classical Flk2+ pathway during aging by initiation of a pathway in parallel to canonical Plt differentiation (**Figure 1I**).

**Table 1.** Percent GFP+ cells in hematopoietic populations from Old FlkSwitch mice.

| Mouse | HSC | MPP | CMP | GMP | MEP | CLP | MkP | Plt | EP | CFU-E | pCFU-E | pGMP | pMEgE | "GMP" | GM | B cell | T cell |
| --- | --- | --- | --- | --- | --- | --- | --- | --- | --- | --- | --- | --- | --- | --- | --- | --- | --- |
| 1 | 1.8 | 96.7 | 86.5 | 86.5 | 92.0 | 100.0 | 42.2 | 48.4 | 96.5 | 86.5 | 88.6 | 90.7 | 85.1 | 91.1 | 96.6 | 98.4 | 98.4 |
| 2 | 0.0 | 94.3 | 81.1 | 88.0 | 84.9 | 100.0 | 38.9 | 48.1 | 90.8 | 85.3 | 79.2 | 83.6 | 80.0 | 89.2 | 86.1 | 95.5 | 94.0 |
| 3 | 1.9 | 98.0 | 81.8 | 90.9 | 86.1 | 100.0 | 33.6 | 43.9 | 85.7 | 87.2 | 85.4 | 84.6 | 79.5 | 92.0 | 89.0 | 88.9 | 94.6 |
| 4 | 3.8 | 91.7 | 82.0 | 93.8 | 82.3 | 93.0 | 29.6 | 35.9 | 96.7 | 93.5 | 88.4 | 93.7 | 91.9 | 97.3 | 95.6 | 96.0 | 98.8 |
| 5 | 2.9 | 98.5 | 87.8 | 94.5 | 87.7 | 90.1 | 32.0 | 40.3 | 91.8 | 90.0 | 86.8 | 86.6 | 88.0 | 90.3 | 92.1 | 94.9 | 95.9 |
| 6 | 2.1 | 100.0 | 84.4 | 96.3 | 89.8 | 100.0 | 35.5 | 46.9 | 97.1 | 93.4 | 87.9 | 89.6 | 89.9 | 94.6 | 97.6 | 92.8 | 96.2 |
| 7 | 1.2 | 91.3 | 90.0 | 94.3 | 89.4 | 95.9 | 58.2 | 41.2 | 90.3 | 90.4 | 90.8 | 90.8 | 89.1 | 94.8 | 87.5 | 97.7 | 99.3 |
| 8 | 3.6 | 84.2 | 83.9 | 89.2 | 83.7 | 90.0 | 24.2 | 33.6 | 88.8 | 86.8 | 80.2 | 86.4 | 83.6 | 91.4 | 90.3 | 98.0 | 98.2 |
| 9 | 5.6 | 86.7 | 94.4 | 98.3 | 94.7 | 100.0 | 37.6 | 44.1 | 96.5 | 97.3 | 93.5 | 96.4 | 87.9 | 99.1 | 96.7 | 96.9 | 99.2 |
| 10 | 2.0 | 97.7 | 72.0 | 81.9 | 79.0 | 100.0 | 19.8 | 35.5 | 91.5 | 80.8 | 81.1 | 83.7 | 66.3 | 83.6 | 94.2 | 93.8 | 98.6 |

### Young and Old HSCs similarly undergo differentiation through the Flk2+ pathway upon transplantation

Several reports have clearly documented that the BM microenvironment undergo significant changes with aging and that these changes affect HSPC phenotype and function (Ho et al. 2019; Young et al. 2021; Maryanovich et al. 2018; Saçma et al. 2019). To determine whether the Plt-specific pathway was imposed by the aged environment or manifested by heritable changes in aged HSCs, we performed heterochronic and isochronic transplantation experiments of young or old HSCs from FlkSwitch mice and monitored the differentiation paths by tracking Tom/GFP fluorescence ratios of donor-derived mature cells in the PB (**Figure 2A and C**). As we have previously shown, the vast majority of circulating cells derived from the transplanted Tom+ yHSCs were GFP+ (>90% of B and T cells were GFP+, and Plts and GMs plateaued at ∼60%) after transplantation into young hosts (Y-Y) (Boyer et al. 2011). To determine whether oHSCs deviate from this baseline Tom/GFP ratio, and therefore retain the age-specific Plt differentiation potential, we transplanted old FlkSwitch HSCs into young recipients. Interestingly, oHSCs gave rise to Plts, GMs, B, and T cells with similar Tom/GFP ratios (O-Y) as yHSCs (Y-Y) in young recipients, indicating that transplanted oHSCs did not retain their divergent Plt specification in the young niche (**Figure 2B**). Similar Tom/GFP patterns were obtained when either yHSCs (Y-O) or oHSCs (O-O) were transplanted into Old recipients (**Figure 2D**). Thus, the aged niche did not appear to induce age-specific differentiation of transplanted yHSCs, and neither hetero-nor isochronic transplantation of oHSCs recapitulated the divergent Plt differentiation path observed in situ. Instead, transplantation appeared to reset oHSCs into a youth-like differentiation path, regardless of recipient age and despite reduced overall repopulation capacity (**Figure S4**) (Poscablo et al. 2021). These experiments demonstrate that the differentiation paths of both yHSCs and oHSCs are relatively unaffected by the age of the recipient environment in adoptive transfer experiments.

**Figure 2.**
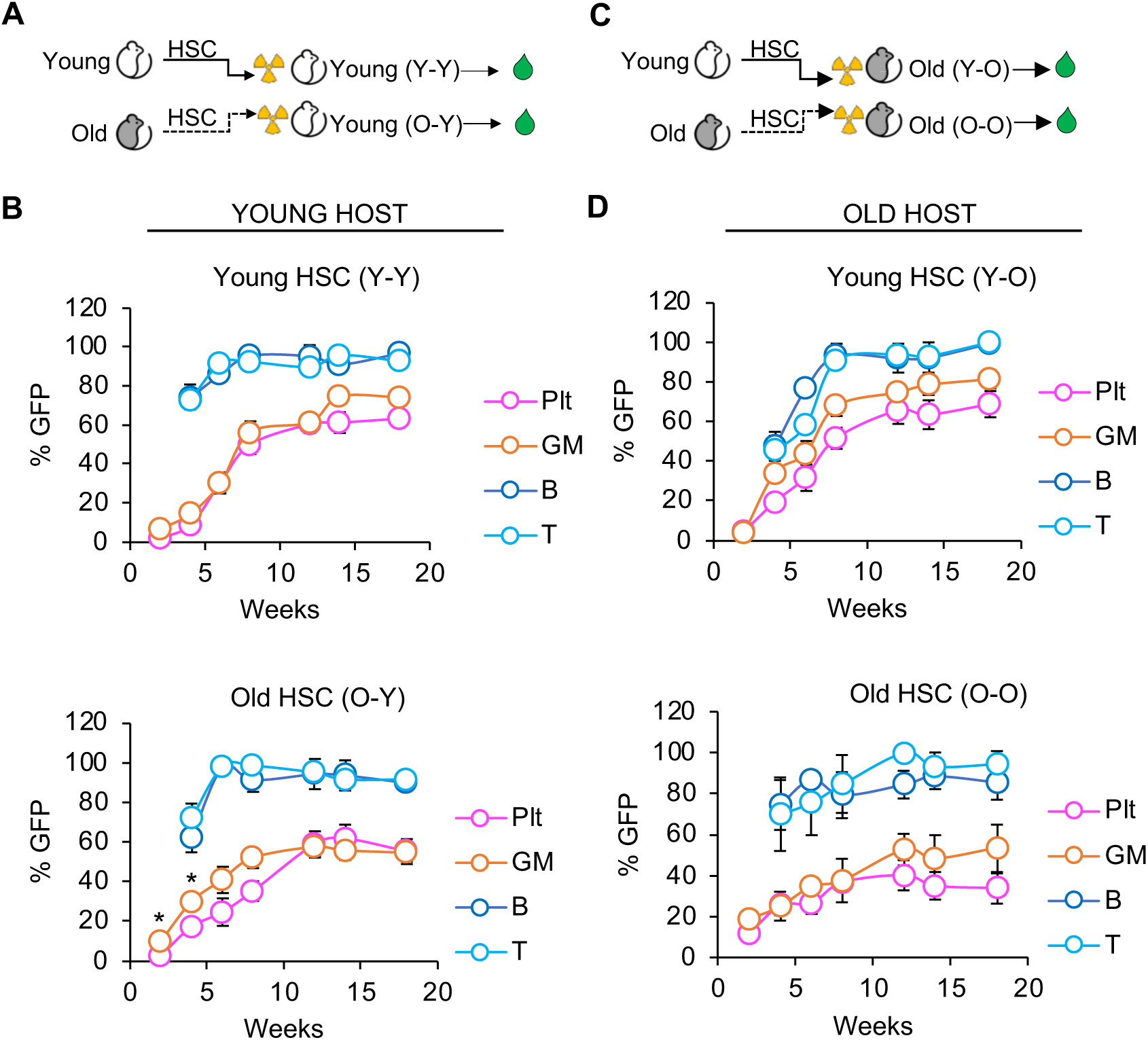
Old HSCs did not retain the age-specific Plt differentiation pathway upon transplantation. **A and C.** Schematic of heterochronic and isochronic transplantation of HSCs from FlkSwitch into conditioned, non-fluorescent mice. Peripheral blood was monitored for GFP/Tom+ fluorescence of donor-derived mature cells, presented as %GFP in recipient mice. **B and D.** Percentage of GFP+ donor-derived cells were equivalent in young (B) or old (D) recipients of transplanted young or old HSCs for >16 weeks post-transplant. Data represent mean ± SEM of 5 independent experiments with n= 15 Y-Y mice, n= 21 O-Y mice, n= 13 Y-O mice, n= 13 O-O mice. Statistics: unpaired two-tailed t-test between Plts and GMs. *p<0.05. Y-Y, Young HSCs transplanted into young recipients; O-Y, Old HSCs transplanted into young recipients; Y-O, Young HSCs transplanted into old recipients; O-O, Old HSCs transplanted into old recipients.

### Aging induces divergent transcriptome alterations in age-specific MkPs

To investigate the molecular programs of age-induced MkP heterogeneity, we compared the transcriptomes of the three distinct murine MkP populations: GFP+ yMkP, GFP+ oMkP, and Tom+ oMkP. GFP+ MkPs remained relatively similar with age, distinguished by only 90 differentially expressed genes (DEGs) (**Figure 3A**). This indicates a remarkable resilience of GFP+ oMkPs to age-induced changes known to be characteristic of the BM environment (Ho et al. 2019; Young et al. 2021; Maryanovich et al. 2018; Saçma et al. 2019). In contrast, Tom+ oMkPs were substantially different from both GFP+ yMkPs (1,150 DEGs) and oMkPs (200 DEGs). Interestingly, a number of genes associated with HSC-selective expression were found to be increased in expression in the Tom+ oMkPs compared to GFP+ oMkPs, such as *Mllt3, Fhl1, Plscr2, Nupr1, Cdkn1c,* and *Hoxb5* (**Figure 3B**) (Calvanese et al. 2019; Ivanova et al. 2002; Wang et al. 2022; Matsumoto et al. 2011; Chen et al. 2016; Forsberg et al. 2010). We corroborated our gene expression data by using flow cytometry to demonstrate that some mRNA changes resulted in differential expression of cell-surface proteins of one representative upregulated (CD105) and downregulated (CD119) in Tom+ vs GFP+ oMkPs (**Figure 3C**). These data demonstrate that Tom+ oMkPs represent a novel age-selective hematopoietic progenitor with an age-unique molecular profile. To investigate the mechanisms by which aging induces the divergent Plt differentiation pathway, we compared expression profiles of young and old MkPs to oHSCs. Interestingly, compared to GFP+ yMkPs and GFP+ oMkPs, Tom+ oMkPs were transcriptionally more similar to oHSCs (**Figure 3D, Figure S5A)**. Hierarchical clustering analysis of gene expression patterns also confirmed distinct association of the three MkP population and oHSCs and revealed a subset of genes that are highly expressed in both Tom+ oMkP and oHSCs, but not GFP+ MkPs (**Figure 3E).** Tom+ oMkP also clustered closest to oHSCs when we compared our RNAseq data with HSC- and MkP-specific genesets available from Gene Expression Commons (GEXC) (**Figure S5B**) (Seita et al. 2012). Together, these demonstrated that oHSCs were transcriptionally closer to Tom+ oMkPs compared to old and young GFP+ MkPs, supporting the shortcut pathway from oHSCs into Tom+ oMkPs identified by lineage tracing.

**Figure 3.**
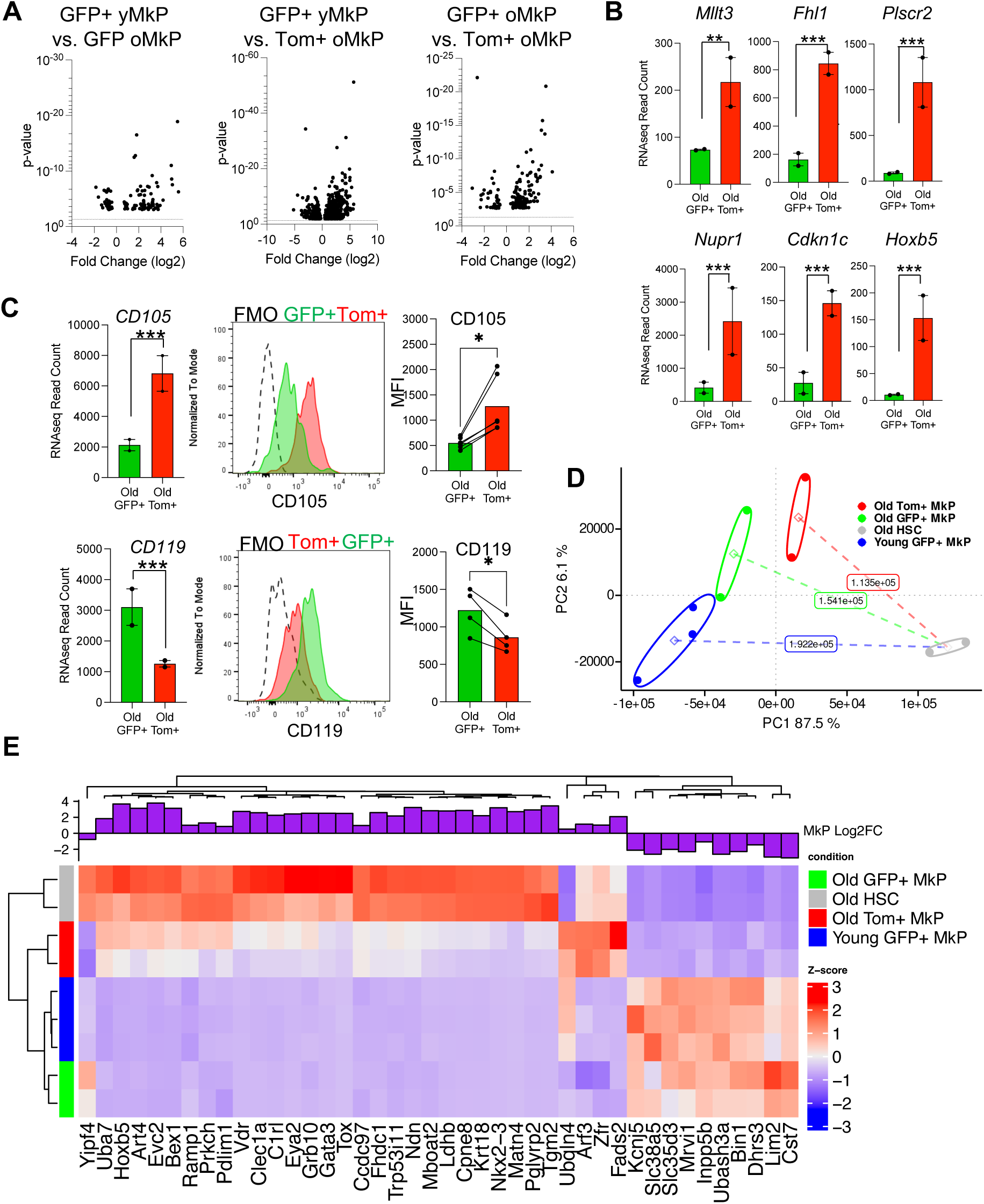
RNA-Seq analysis revealed a distinct molecular profile of Tom+ MkPs. **A.** Volcano plots showing differentially expressed genes (DEGs) expressed between GFP+ yMkPs vs GFP+ oMKPs, between GFP+ yMkPs vs Tom+ oMkPs, and between GFP+ oMkPs vs Tom+ oMkPs. Dotted lines indicate where p value = 0.05. **B. Tom**+ oMkPs highly express genes associated with HSC function. mRNA levels of specific DEGs by RNAseq read count in Tom+ oMkPs compared to GFP+ yMkPs. **P<0.005, ***P<0.0001. **C.** Differential cell surface protein expression of CD105 and CD119 by the two populations of old MkPs. Results from RNAseq analysis were tested by differential flow cytometry for CD105 and CD119 on GFP+ oMkPs and Tom+ oMKPs. MFI Data represent 3 independent experiments with n=4 young, n=4 old mice. Statistics: paired t-test. *P<0.05. RNAseq analysis as in **B.** **D.** Tom+ oMkPs are located closest to oHSCs in transcriptional space. Principal Components 1 and 2 capture 93.6% of the total transcriptional variance across MkPs and HSCs, demonstrating that Tom+ oMkPs are most similar to oHSCs in PC1 and according to Euclidean distance calculated from centroids (opaque diamonds) for each cell type. **E.** Kmeans-clustered heatmap of gene expression Z-score for significantly differentially expressed genes between Tom+ oMkPs (up) and GFP MkPs (down) demonstrates shared transcriptional signal in oHSCs and Tom+ oMkPs that is diminished or absent in other MkP populations. yMkP, young MkP; oMkP, old MkP; oHSC, old HSC

### Age-specific MkPs are functionally enhanced compared to both the canonically-derived coexisting Old MkPs and to Young MkPs

Our observations that transplanted Tom+ yHSCs and oHSCs exhibit similar floxing efficiency and the absence of a uniquely divergent Plt differentiation pathway from transplanted HSCs point to oMkPs as putative major perpetuators of the age-specific Plt pathway. Consistent with functional divergence of progenitors upon aging, we recently demonstrated that the bulk population of oMkPs have a remarkable expansion capacity compared to yMkPs (Poscablo et al. 2021). While these alterations could logically have been attributable simply to the process of either cell-intrinsic aging or to an aging phenotype imposed by the environment, our RNAseq data where GFP+ MkPs were more similar throughout aging than to Tom+ MkPs that co-exist with GFP+ MkPs in the aged environment prompted us to consider alternative explanations. To investigate potential functional consequences of age-specific megakaryopoiesis, we first compared the in vitro expansion capacity of GFP+ yMkPs, GFP+ oMkPs, and Tom+ oMkPs (**Figure 4A**). Whereas the GFP+ yMkPs and GFP+ oMkPs were indistinguishable in their low capacity to expand in culture, Tom+ oMkPs significantly expanded compared to both GFP+ MkP populations (**Figure 4B**). These observations prompted our assessment of the three MkP populations by transplantation (**Figure 4C**). We previously demonstrated that the bulk population of oMkPs robustly reconstitute Plts, therefore we were surprised to find that the GFP+ fraction of oMkPs minimally contributed to Plt donor-chimerism and at no greater capacity than yGFP+ MkPs (reaching 1-6% donor contribution) (**Figure 4D**). In direct contrast to both GFP+ MkP populations, the Tom+ oMkPs were remarkably capable of robust, but transient, reconstitution of Plts (**Figure 4D**). Consistent with our reported results, all three MkP populations lacked the capacity to reconstitute B and T cells, while some erythroid and GM chimerism was observed; this was primarily by Tom+ oMkPs (**Figure S6A)** (Poscablo et al. 2021). Similar to a recent study (Morcos et al. 2022), we found that yMkPs can be phenotypically distinguished by CD48 expression, with detection of a small population of CD48-MkPs in young, unperturbed mice (**Figure S6B**). Importantly, however, the CD48+ and CD48-yMkP populations were functionally indistinguishable from each other by in vitro analysis (**Figure S6C**) and did not have higher Plt reconstitution capacity upon transplantation (**Figure S6D**), but did produce significantly more erythroid cells (**Figure S6E**). These data from WT mice are consistent with the FlkSwitch model in that the vast majority of MkPs in Y mice can be viewed as one population under native conditions. Collectively, the surprising gain of functional capacity upon aging that directly contrast the behavior of aged HSCs appears to be harbored entirely within the age-specific Tom+ MkPs (Poscablo et al. 2021). This enhanced capacity can be mechanistically explained by their specification from HSCs via shortcut differentiation, with inheritance of sustained expression of stem cell-promoting genes such as *Hoxb5*, *Mllt3*, *c-Kit*, and others (**Figure 3B**) that, collectively, contribute to the observed increase in lineage potential and engraftment capacity. The superior ability of age-specific MkPs to expand, engraft, and reconstitute Plts compared to the co-existing canonical MkPs suggests a selective and pathway-specific consequence of aging. Moreover, the functional similarities in GFP+ yMkPs and GFP+ oMkPs further support a model where the canonical, Flk2+ differentiation path tempers major age-imposed changes to molecular and functional properties throughout life.

**Figure 4.**
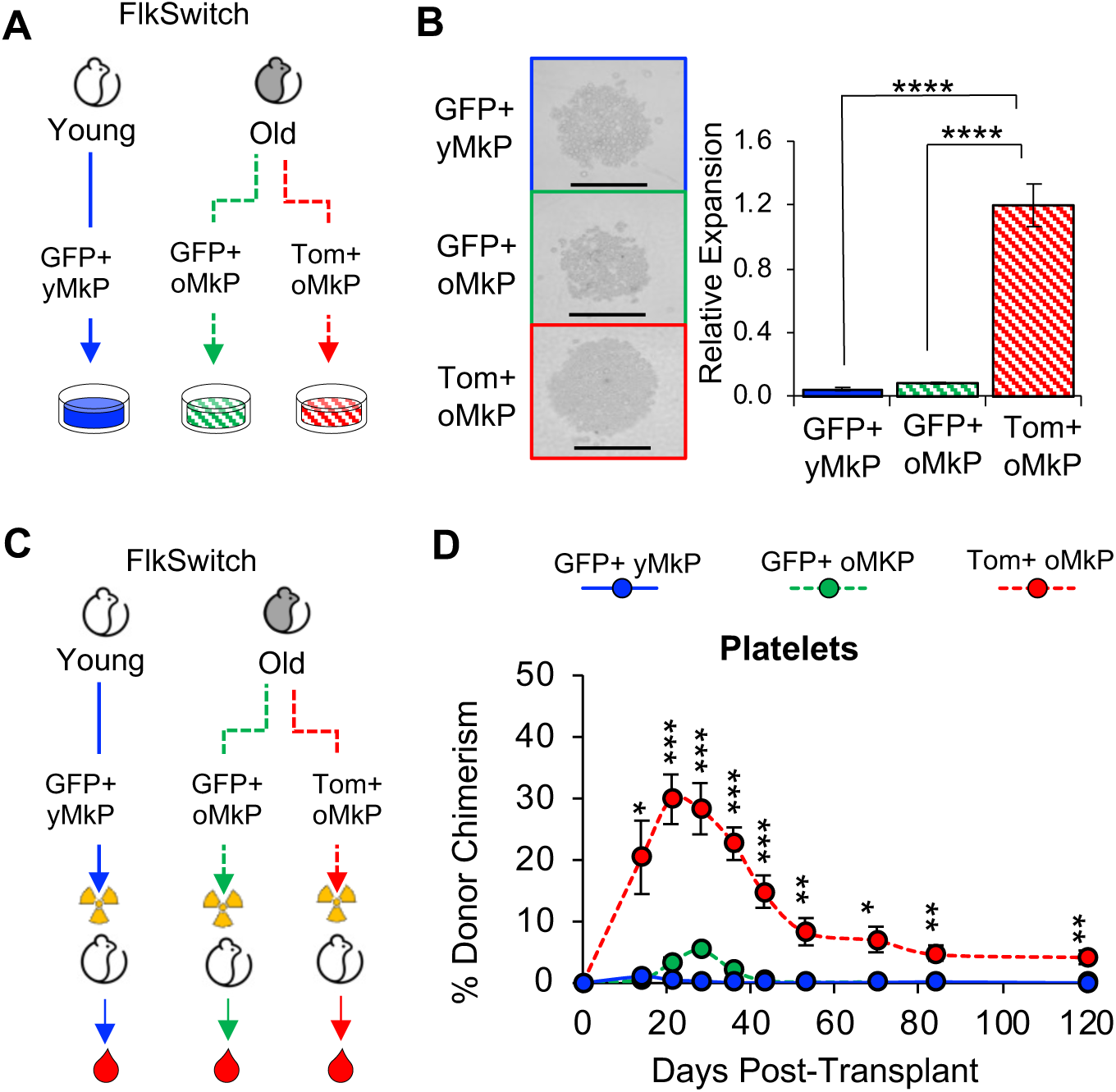
Tom+ MkPs from old FlkSwitch mice have greater expansion and platelet reconstitution potential compared to GFP+ Young or Old MkPs. **A.** Schematic of in vitro expansion assay of MkPs from young or old FlkSwitch mice. 1000 GFP+ yMkPs, GFP+ oMkPs, or Tom+ oMkPs were isolated by FACS and plated per well. After 3 days of expansion, cells were quantified by flow cytometry. **B.** Tom+ oMkPs displayed greater expansion capacity in vitro compared to both GFP+ yMkPs and GFP+ oMkPs. Left: Representative images of MkP in vitro cultures at day 3. Right: Quantification of cell expansion revealed significantly greater number of cells from Tom+ MkPs compared to both GFP+ MkPs. Data represent mean ± SEM of 3 independent experiments, n = 9 GFP+ yMkP wells, n = 9 GFP+ oMkP wells and n = 17 Tom+ oMkP wells. Statistics: one-way anova and Tukey post hoc test. **** p < 0.001 **C.** Schematic of MkP transplantation from young or old FlkSwitch mice. 22,000 GFP+ yMkPs, GFP+ oMkPs, or Tom+ oMkPs were isolated by FACS and transplanted into young WT recipient mice. Peripheral blood analysis by flow cytometry was done to monitor repopulation of mature cells. **D.** Old Tom+ MkPs demonstrated greater contribution to platelets in the recipient mice compared to both GFP+ yMkPs and GFP+ oMkPs. Analysis of donor-derived Plts in peripheral blood of recipients presented as percent donor chimerism. Data represent mean ± SEM of 3 independent experiments, n = 6 GFP+ yMkP recipients, n = 4 GFP+ oMkP recipients, and n = 13 Tom+ oMkP recipients. Statistics: unpaired two-tailed t-test. T-tests between GFP+ yMkP and GFP+ oMkP were not statistically significant. T-tests between GFP+ oMkP and Tom+ oMkP *p<0.05, **p<0.005, ***p<0.0005. yMkP, young MkP; oMkP, Old MkP

### Age-specific platelets participate in exacerbated clot formation upon vascular injury

The predominant function of megakaryopoiesis is to generate Plts that are essential for hemostasis. Age-related dysregulation of both Plt numbers and activity poses tremendous health risks for aging humans. Here, we show that the well-documented significant increase in Plt counts in aging mice is due to an additive accumulation of Tom+ Plts to the canonical GFP+ Plts, while changes to the absolute number of other circulating mature cells result from GFP+ differentiation pathways (**Figure 5A)** (Culmer et al. 2013; Poscablo et al. 2021; Davizon-Castillo et al. 2019). Remarkably, despite being specified by two molecularly and functionally distinct paths, immunophenotyping of the Tom+ oPlts revealed that they uniformly expressed well-established Plt surface proteins, including CD41, CD9, CD42a, and CD42b (**Figure 5B**). To assess Plt functionality, we evaluated the role of Tom+ and GFP+ Plts in clot formation in vivo using an intravital microscopy laser ablation model of hemostasis in real time (**Figure 5C)** (Tourdot et al. 2017; Adili et al. 2017; Reheman et al. 2009). We introduced laser-induced ablation injuries to cremaster arteriole walls of young and old FlkSwitch mice to initiate and directly visualize thrombus formation, then quantified the accumulation of Tom+ and GFP+ thrombus constituents in real time. Vascular injury to young FlkSwitch mice resulted in the formation of small, unstable thrombi composed of only GFP+ cells (**Figure 5D, top panel, Figure S7, Movie S1**). In contrast, immediately after rupture of the vascular walls in old FlkSwitch mice, we observed the formation of dramatically large, stable thrombi at the site of injury, with participation of both GFP+ and Tom+ Plts (**Figure 5D, bottom panel, Figure S7, Movie S2)**. Larger thrombus formation in the old FlkSwitch model were substantiated by significantly greater fluorescence exhibited by GFP+ and Tom+ cells within the clots of the old FlkSwitch mice compared to young FlkSwitch mice (**Figure 5E**). Additionally, vascular insult also resulted in greater accumulation of fibrin within the clot of old mice compared to young mice, indicative of more stable thrombi (**Figure 5F**). These experiments uncovered that laser-induced clot formation is drastically amplified in old mice, and that both Tom+ and GFP+ Plts contribute to the enlarged thrombi.

**Figure 5.**
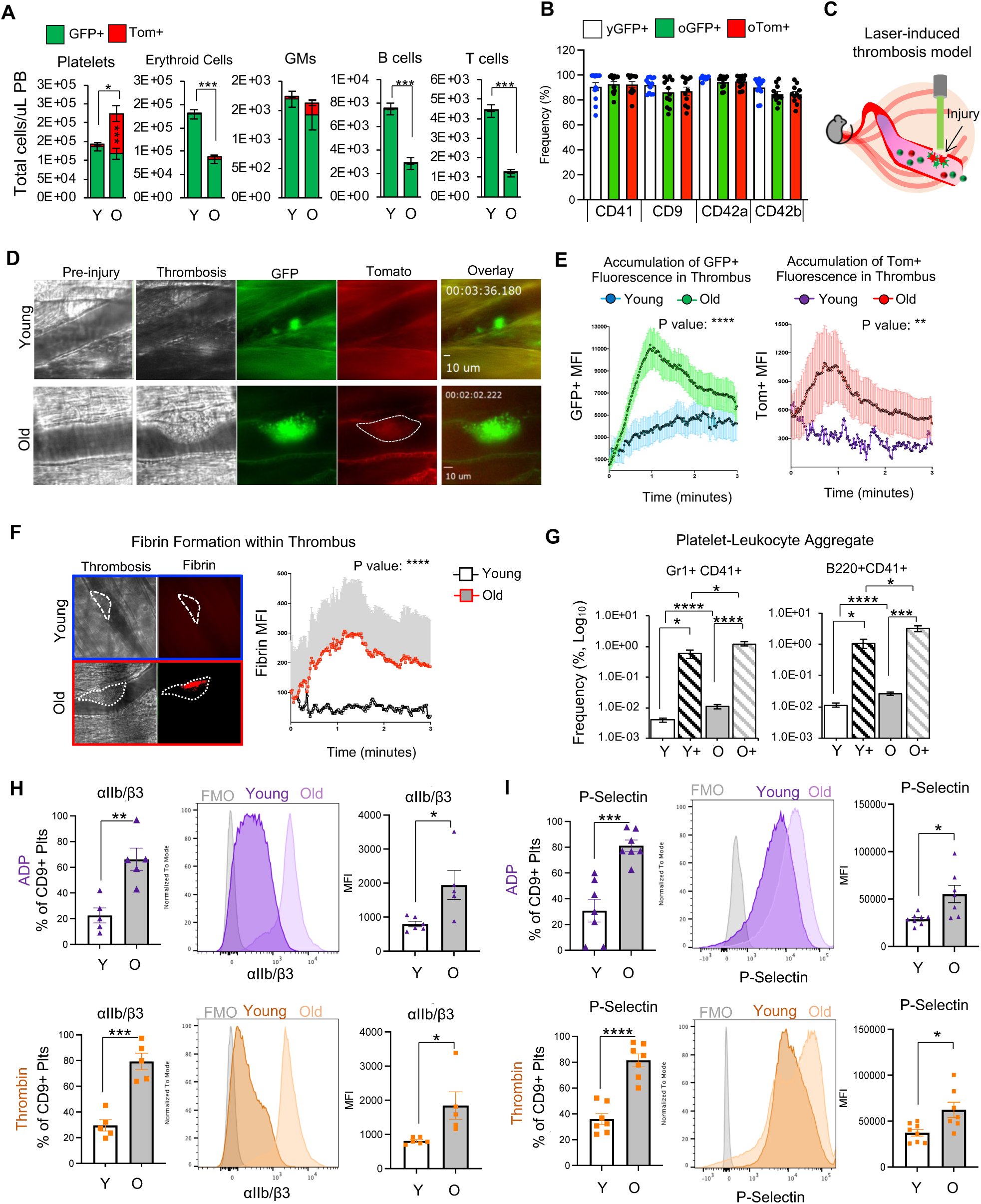
The age-specific Plt pathway contributes to Plt hyperactivity in old FlkSwitch mice. **A.** Aging leads to thrombocytosis due to accumulation of Tom+ Plts. Absolute quantification of circulating cells presented as total cells/microliter of PB. While changes to the absolute number of other mature cells in the PB result from cells derived via GFP+ differentiation pathways, the numerical increase in Plts is a consequence of the Tom+ differentiation path in aged FlkSwitch mice. Statistics: unpaired t-test. *P<0.05, **P<0.005, ***P<0.0005. T-tests between Tom+ yPlts and Tom+ oPlts: ***P<0.0005. T-tests between all other Tom+ Y and Tom+ O cells were not significant. **B.** Tom+ oPlts express traditional Plt markers. Frequency of Plts expressing known Plt surface markers: CD41, CD9, CD42a, CD42b (GPIb**α)**. yGFP, young GFP+ Plts; oGFP, old GFP+ Plts; oTom+, old Tom+ Plts. **C.** Schematic of the laser-induced thrombosis model to monitor thrombus formation upon vascular injury. **D.** Tom+ oPlts contribute to excessive thrombus formation in old mice. Representative images of clot formation in Young (top) and Old (bottom) FlkSwitch mice displaying participation of GFP+ and Tom+ Plt in thrombi. Also see Supplemental Movies. **E.** Tom+ and GFP+ cells are major contributors to greater thrombus formation in old FlkSwitch mice, while the smaller thrombi in young FlkSwitch mice consist exclusively of GFP+ cells. Dynamics of clot formation at time points post vascular injury were quantified by MFI of thrombi, comparing GFP+ (left) or Tom+ (right) accumulation in young or old FlkSwitch mice. Data represent mean ± SEM of 3 independent experiments with n = 3 Y, and n =3 O mice, with 9-12 injuries per mouse. Statistics: paired t-test. ****P<0.0001, **P<0.005 **F.** Fibrin content increased in thrombi in old mice. Dynamics of fibrin formation in thrombi at time points post vascular injury were analyzed by change in fluorescent intensity conferred by A647-conjugated anti-fibrin antibodies. Representative images of fibrin formation (left) and quantification of fibrin MFI within thrombi (right) in young or old FlkSwitch mice. Data represent 3 independent experiments with n = 3 young, and n =3 old mice. Statistics: paired t-test. ****P<0.0001 **G.** Platelet-leukocyte aggregate formation is greater in old blood compared to young blood. Quantification of mature myeloid cell aggregation with Plts (Gr1+CD41+, left) and B-cell aggregation with Plts (B220+CD41+, right) with and without thrombin-mediated activation (0.1 units/mL). Y, young without thrombin; Y+, young with thrombin; O, old without thrombin; O+, old with thrombin. Data represent 3 independent experiments with n = 7 Y mice, n = 8 Y+, mice n= 22 O mice, n= 8 O+ mice. Statistics: one-way anova and Tukey post hoc test. *P<0.05, ***P<0.0005, ****P<0.0005 **H.** oPlts demonstrated higher activation of integrin αIIb/β3 upon stimulation by ADP (10 µM) and thrombin (0.1 U/mL). Quantification of αIIb/β3+ Plt frequency (left of flow cytometry histogram) and αIIb/β3 MFI (right) within CD9+ Plts. Data represent 3 independent experiments with n = 5 Y mice, n = 5 O mice. Statistics: unpaired t-test. *P<0.05, **P<0.005, ***P<0.0005 **I.** oPlts demonstrated higher P-Selectin surface display upon stimulation by ADP (10 µM) and thrombin (0.1 U/mL). Quantification of P-Selectin+ Plt frequency (left) and P-Selectin MFI (right) within CD9+ Plts. Data represent 3 independent experiments with n = 7 Y mice, n = 7 O mice. Statistics: unpaired t-test. *P<0.05, ***P<0.0005, ****P<0.0005

The hyperreactivity of Plts from old mice was also evident by Plt-leukocyte aggregation (PLA) assays that quantify mature myeloid and B-cell interaction with Plts. Circulating PLAs appeared at significantly higher frequencies in old compared young blood (**Figure 5G, Figure S8**). Upon in vitro stimulation by thrombin, we also observed a greater elevation in PLA formation in old compared to young blood (**Figure 5G, Figure S8**). Thus, PLA formation is elevated in old blood under both basal and stimulated conditions, further emphasizing the potent thrombotic response in aged blood.

We then sought to measure the activation of oPlts by which hyperactive Plts could drive enhanced clot formation in aged blood. Using flow cytometry, we examined the expression of cell-surface proteins that are critical for Plt activation. Upon stimulation, the integrin αIIb/β3 switches from low affinity to high affinity for its ligand (fibrinogen) to promote adhesion and thrombus growth (Huang et al. 2019). Likewise, P-Selectin is translocated from intracellular granules to the external membrane in activated Plts (Merten & Thiagarajan 2000; Ivanov et al. 2019). In vitro analysis of Plt activation revealed conformational change and elevated expression of αIIb/β3 and P-selectin, respectively, on oPlts compared to yPlts upon both adenosine diphosphate (ADP, **Figure 5H**) or thrombin (**Figure 5I**) stimulation. Thus, αIIb/β3 and P-Selectin expression are consistent with hyperreactivity of old Plts in response to stimulatory agonists such as thrombin and ADP. Together, these data suggest that the two Plt populations in aged mice contribute to physiological alterations in hemostasis by enhancing Plt participation in clot formation.

### Age-unique MkPs rapidly restore acutely depleted Plts

HSPCs are amazingly responsive to produce mature cells when provoked by stress, including inflammatory stimuli that trigger megakaryopoiesis (Essers et al. 2009; Bogeska et al. 2022; Haas et al. 2015). Therefore, the drastic increase in thrombocytosis-fueled Plt pathologies and the remarkable Plt reconstitution capacity of age-specific MkPs led us to hypothesize that the age-specific pathway may also harbor highly potent physiological control of Plt homeostasis. To directly test whether the age-specific MkPs endow O mice with enhanced Plt restoration capacity, we elicited acute thrombocytopenia in Y and O FlkSwitch mice and measured Plt-differentiation kinetics during Plt recovery. Plt-specific stress was induced by subjecting Y and O FlkSwitch mice to a single anti-Plt antibody injection (**Figure 6A**). As expected (Bergmeier et al. 2000; Salzmann et al. 2020; Nieswandt et al. 2000), this led to extremely rapid and robust depletion of circulating Plts in Y mice, with gradual recovery starting ∼4 days (∼96 hours, **Figure 6B**). Similarly, Plt counts also dropped drastically in O mice; notably, however, Plt numbers were restored more rapidly compared to Y mice (**Figure 6B**). Analysis of the Tom:GFP ratio of Plts suggested that a significantly greater proportion of the newly produced Plts were derived via the shortcut pathway (**Figure 6C,D**). The Plt-challenge also provoked alterations to BM cellularity at 24 hrs post-Plt depletion, with significant and trending reduction of MkPs the Y mice and O mice, respectively, while no major changes were observed in HSC and MPP cellularity (**Figure 6E,F**). To determine if the Tom+ MkPs may have a competitive advantage to respond to the Plt-challenge, we quantified in vivo 5-ethynyl-2′-deoxyuridine (EdU) incorporation upon Plt depletion. The challenge elicited an overall increase in proliferation of both HSCs and MkPs, while MPPs were unaltered compared to steady-state (**Figure 6G, H**). Importantly, in O mice, consistently marked increase in proliferation was exhibited by Tom+, but not GFP+, oMkPs. The hyper-responsiveness of functionally enhanced age-specific Tom+ oMkPs is consistent with our observations of their uniquely profound reconstitution capacity in transplantation (**Figure 4**). Taken together, these data suggest that HSCs and MkPs exhibit a dynamic response to rebuild the circulating Plt supply with age-specific shift in cellular mechanisms. Compared to GFP+ oMkPs, the age-specific Tom+ oMkPs serve a dominate role as first responders in the rapid rescue of acute thrombocytopenia.

**Figure 6.**
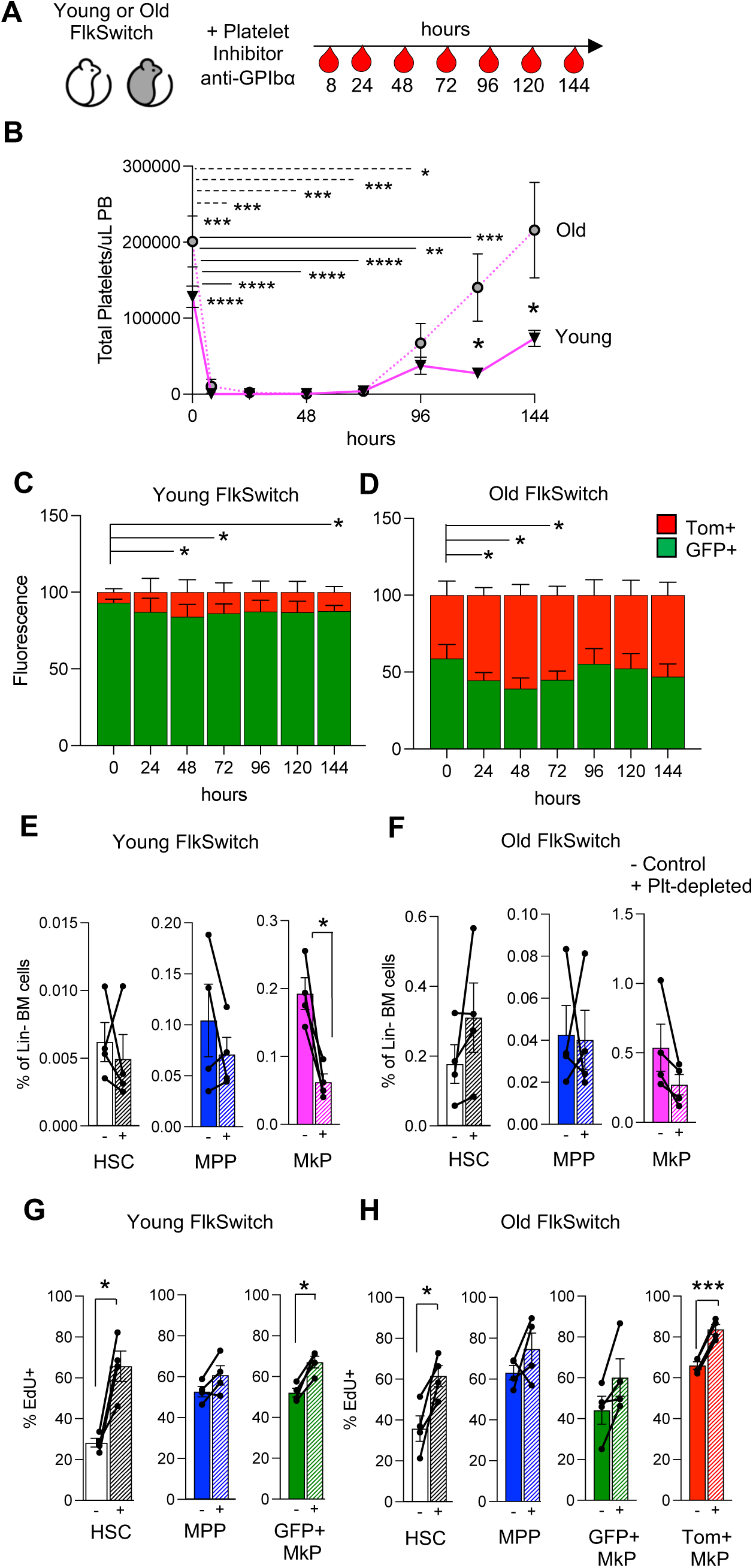
Age-specific MkPs serve as first responders to acute Plt depletion. **A.** Schematic of antibody-mediated Plt depletion and subsequent time-course analyses of cellular response. **B.** A single injection of anti-GPIbα antibodies led to rapid and robust Plt depletion followed by gradual restoration of circulating Plt numbers. Y and O FlkSwitch mice were treated as indicated in panel **A**. Plt numbers in peripheral blood (PB) are indicated at each time point. **C-D.** Analysis of Tom:GFP Plts demonstrated reduction in floxing in Y (C) and O (D) mice during Plt restoration. Data in B-D represent mean±SEM of 4 independent experiments with n = 6 Y mice, n = 4 O mice. Statistics: unpaired t-test. *P<0.05, **P<0.005, ***P<0.0005, ****P<0.00005. **E-F.** Quantification of BM cellularity 24 hours post anti-GPIbα-mediated Plt depletion revealed a significant decrease in frequencies of yMkPs, while yHSCs and yMPPs were unaltered (E). A similar pattern was observed in O mice, albeit an insignificant trend in oMkP cellularity upon Plt-depletion (F). **G-H**. Plt depletion selectively induced MkP proliferation. Short-term in vivo EdU incorporation revealed that HSCs and MkPs, but not MPPs, in Y and O mice respond to Plt depletion, with Tom+ oMkPs dominating the response in O mice (H). Proliferation rates in control and Plt-depleted mice pulsed for 24 hours with EdU, with or without simultaneous administration of anti-GPIbα antibodies. Data in E-H represent mean ± SEM of 4 independent experiments with n = 4 Y mice, n = 4 O mice. Statistics: paired t-test. *P<0.05, ***P<0.0005.

## DISCUSSION

### Discovery of a Plt-specific HSC differentiation path in aging

Here, we utilize the unmanipulated, native hematopoietic system of the FlkSwitch mouse to investigate HSC lineage output from young adulthood into aging. Our study revealed that the Plt lineage uniquely deviates from the classical Flk2+ pathway during aging, with Tom+ HSCs directly differentiating into accumulating Tom+ MkPs that generate Tom+ Plts by bypassing the co-existing canonical differentiation cascade of Flk2Cre-marked GFP+ progenitors. Consequently, the age-specific differentiation pathway leads to the production of a second Plt population that causes steady state-thrombocytosis, exacerbated injury-induced thrombosis, and rapid thrombopoiesis from a stress-responsive pool of aging-unique MkPs in aged mice.

### Native versus stress-induced megakaryopoiesis

Functional and molecular heterogeneity of HSPCs has been well documented (Sanjuan-Pla et al. 2013b; Shin et al. 2014; Grinenko et al. 2014; Pietras et al. 2015; Yamamoto et al. 2013), with stress and aging implicated in selectively promoting megakaryopoiesis(Mitchell et al. 2023; Haas et al. 2015). Previous reports have concluded that HSCs may be primed for rapid and unilineage differentiation into platelet-committed cells (Rodriguez-Fraticelli et al. 2018; Roch et al. 2015; Sanjuan-Pla et al. 2013a; Yamamoto et al. 2013; Haas et al. 2015). Here, we unequivocally demonstrate that a bypass path to Plt production exists in aging mice (**Figure 1**). In contrast, a direct HSC-MkP path in young mice is incompatible with lineage tracing data by us and others (**Figure 1**) (Boyer et al. 2011; Buza-Vidas et al. 2011). This age-dependency is further supported by the absence of functionally enhanced MkPs in young mice (**Figure 4 and S6**). Potential reasons for perceived detection of direct HSC-MkP differentiation in other studies include the use of in vitro differentiation systems; inferring differentiation from transcriptional priming; reliance on assumptions incorporated into complex mathematical models; and use of inducible systems that cause HSC proliferation and potential lineage bias. Indeed, both pIC and tamoxifen, two of the most commonly used agents of inducible lineage tracing systems, selectively induces HSC cycling and differentiation into the megakaryocyte lineage (Sánchez-Aguilera et al. 2014; Sánchez-Aguilera & Méndez-Ferrer 2016; Haas et al. 2015). The strength of our model lies in its simplicity: we simply enumerate functionally defined cells as Tom+ or GFP+ without inducing perturbations and without making assumptions or inferences by mathematical modeling (**Figure 1**). Cells that are GFP+ have (at some point) expressed sufficient levels of Flk2-driven Cre recombinase; cells that remain Tom+ have not. Under native conditions in young adult mice, HSCs differentiate via canonical progenitor cells; aging may serve as a stress mechanism that progressively activates a latent, but primed, transcriptional program that ultimately triggers the bypass Plt path (**Figure 1**). Thus, two functionally and molecularly distinct subsets of aged MkPs, independently produced by HSCs via two separate but parallel differentiation paths, mediate Plt production during aging. We showed that thrombocytosis is driven by age-induced heterogeneity of the MkP population (**Figure 5A**), and that an increasing proportion of the Plt homeostatic responsibilities of HSCs appear to be delegated to MkPs upon aging (**Figure 6**), possibly due to the age-related functional decline of HSCs (**Figure S4**). Our prospective isolation of age-specific MkPs pinpoints the cellular mechanism of our recently reported increase in MkP repopulation capacity upon aging (Poscablo et al. 2021; Poscablo & Forsberg 2021) and identifies the age-selective Tom+ MkPs as a potent source of thrombopoiesis.

### Unexpected microenvironment-independent mechanisms of stem and progenitor cell aging

Surprisingly, the canonical GFP+ MkPs maintain youthful properties throughout life, with relatively attenuated expansion and engraftment capacities that are indistinguishable from young GFP+ MkPs. The functional similarities of the young and aged GFP+ MkPs were unexpected, as we detected significant changes in gene expression (**Figure 3**) and because the well-documented changes to the BM environment were expected to alter cell function (Frisch et al. 2019; Stegner et al. 2017; Heazlewood et al. 2023; Zhao et al. 2014; Bruns et al. 2014; Ho et al. 2019). Similar resilience to external factors has recently been reported for HSCs upon attempts to influence HSC aging by several heterochronic strategies (Ho et al. 2021). In stark contrast to the age-resilient GFP+ oMkPs, the Tom+ oMkPs, despite co-existing in the same aged environment as the GFP+ oMkPs, have dramatically enhanced Plt production capacity in vitro and upon transplantation (**Figure 4**) and in response to Plt demand (**Figure 6**). This uncouples the aging microenvironment from MkP functional capacity and demonstrates that MkP properties are influenced to a greater extent by their differentiation path than by the environment of the aged BM. In addition to uncovering hallmarks of megakaryopoiesis, the surprising gain of function of age-specific MkPs confers a new perspective on the potential mechanisms behind the well-established functional decline of oHSCs.

### Controlling age-dependent thrombosis via stem and progenitor cells

While drastic increases in platelet dysregulation and adverse thrombotic events in aging populations have been clear for decades, distinct differentiation paths and entirely novel, age-specific cell populations have not been envisioned as underlying mechanisms of disease susceptibility. The emergence of Plt subpopulations during age-specific ontogeny provides evidence that Plt heterogeneity is a determinant of age-related Plt diseases. Remarkably, functional Plts can be produced via two alternative pathways, one of which is age-dependent and decoupled from erythropoiesis. The production of mature cells via distinct differentiation paths offers a paradigm of stem cell aging that is currently unexplored. Our identification of the cellular origins and mechanisms of age-specific Plts provides compelling therapeutic opportunities for targeting hematopoietic stem cells and megakaryocyte progenitors to control both Plt generation and function throughout life. Therefore, our findings may profoundly impact the millions of elderly people at risk for experiencing adverse thrombotic events.

## Supporting information

Poscablo Supplemental Data

## Acknowledgments

We thank the UCSC Flow Cytometry and stem cell ore facilities, RRIDs SCR_021149, SCR_021353 and SCR_021135. Funding was provided by NIH/NIA R01AG062879 (ECF), NIGMS R25GM058903 (DMP), T32GM8646 (AKW), F31HL151199 (AKW), K12GM139185 (BAM), F99DK131504 (RER) Howard Hughes Medical Institute (DMP), American Heart Association 19PRE34370030 (DMP) Tobacco-Related Disease Research Program (AKW, TC) and the California Institute for Regenerative Medicine CL1-00506; FA1-00617 (UC Santa Cruz).

## Author contributions

DMP and ECF conceived and organized the study and wrote the manuscript, with input from all authors. ECF and MH supervised the work and contributed to the data interpretation. DMP, AKW, BAM, TC, RER, ECF acquired financial resources. DMP, AKW, SSB, BAM, RA, TC, RER, AEB, SWB, and ECF designed and analyzed experiments, and DMP, AKW, SSB, BAM, RA, TC, RER, SD, AEB, and SWB performed experiments.

## Materials & Correspondence

should be directed to C. Forsberg.

## METHODS

### Mouse lines

All animals were housed and bred in the AAALAC accredited vivarium at University of California Santa Cruz and maintained under approved IACUC guidelines. The following mice were utilized for these experiments: C57Bl6 (JAX, cat# 664), aged C57Bl6 (NIH-ROS), UBC-GFP (JAX, cat# 004353), and male FlkSwitch mice (Flt3-Cre x mTmG mice) (Benz et al. 2008; Muzumdar et al. 2007). Young mice were between 8-16 weeks of age and old mice were 20+ months of age, except old transplant recipient mice which were 18+ months of age. All wt C57Bl6 and UBC-GFP mice were randomized based on sex.

### Flow Cytometry

Bone marrow stem and progenitor cell populations and mature cell subsets were prepared and stained as previously described (Smith-Berdan et al. 2019; Rajendiran et al. 2020; Martin et al. 2020; Beaudin et al. 2016; Boyer et al. 2019; Poscablo et al. 2021). Briefly, the long bones (femur and tibia) from mice were isolated, crushed with a mortar and pestle, filtered through a 70μm nylon filter and pelleted by centrifugation to obtain a single cell suspension. Cell labeling was performed on ice in 1X PBS with 5 mM EDTA and 2% serum. HSCs (Lin-/cKit+/Sca1+/Flk2-/Slam+/Tom+) or MkPs (Lin-/cKit+/Sca1-/CD41+/Slam+/Tom+ or GFP+ or Lin-/cKit+/Sca1-/CD41+/Slam+/CD48+ or CD48^lo/-^) from young or old FlkSwitch mice were analyzed from unfractionated samples or isolated from c-Kit-enriched BM with CD117-microbeads (Miltenyi) using a FACS ARIA II (Becton Dickinson, San Jose, CA) as previously described (Poscablo et al. 2021; Boyer et al. 2011; Boyer et al. 2019). Tissue-resident macrophages from brain and lung were analyzed as described (Leung et al. 2019). Brain microglia were analyzed as Live, CD45+/F4/80hi/CD11bhi/Ly6g−/CD11c− and lung alveolar macrophages were analyzed as CD45+F4/80hi/CD11bmid/SiglecF+. Cells with no history of Cre expression were defined as Tom+GFP-(“unfloxed”) cells, whereas cells with current or a history of Cre expression were defined solely by GFP expression, based on our previous demonstration that Tom+GFP+ cells have excised the loxP-flanked Tomato cassette(Beaudin et al. 2016; López et al. 2022; Boyer et al. 2011; Boyer et al. 2012); that Tom+GFP+ and Tom-GFP+ cells are functionally indistinguishable(Beaudin et al. 2016; Boyer et al. 2011); and well-accepted field standards of loxP-stop-loxP-inducible single-color reporters.(Säwen et al. 2018; Chapple et al. 2018; Buza-Vidas et al. 2011; Morcos et al. 2022)

### In vitro MkP Expansion

MkPs were sorted as described above (1000 per well from FlkSwitch mice or 2000 per cell from wt mice in 96-well U-bottom tissue culture plates) were cultured for 3 days in 200μl/well containing IMDM medium (Fisher) supplemented with 10% FBS, 20ng/ml rmTPO, 20ng/ml rmIL-6, 50ng/ml of rmSCF, and 5ng/ml rmIL-3 (cytokines from Peprotech), 1X Primocin (Invivogen), and 1X non-essential amino acids (Gibco). On day of analysis, a known number of APC-labeled spherobeads (BD Bioscience) were added. Cells were stained as described above and data was collected on either a LSR II, FACSAria IIu (Beckton Dickinson) or CytoFlex LX (Beckman Coulter). Analysis was performed in FlowJo V9 or V10 (Beckton Dickinson). Cell expansion was calculated based on the number of beads recovered per beads added per well, as previously described (Poscablo et al. 2021).

### Transplantation Reconstitution assays

Reconstitution assays of FlkSwitch cells were performed by transplanting HSCs (200 per recipient) or MkPs (22,000 per recipient) from young or old mice into congenic, sublethally irradiated C57Bl6 mice via retro-orbital intravenous transplant as previously described (Cool et al. 2020; Worthington et al. 2022; Poscablo et al. 2021). Reconstitution of wt MkPs was performed by transplanting 22,000 CD48+ or CD48^lo/-^ MkPs from young male and female mice into sublethally irradiated (5 Gy) male and female UBC-GFP hosts via retro-orbital intravenous injection. 1X HBSS (Gibco) devoid of cells was instead injected for the Sham controls. Following transplantation, mice were bled via the tail vein at the indicated time points. Donor chimerism of Plts (single, CD11b- Gr1- CD3- B220- Ter119- CD41+) and erythroid cells (single, CD11b- Gr1- CD3- B220- Ter119+ CD41-) was determined on the whole blood fraction. Donor chimerism of B cells (single, live, CD11b- Gr1- CD3-B220+), T cells (single, live, CD11b- Gr1- B220- CD3+), and GMs (single, live, CD3- B220- CD11b+ Gr1+) were quantified following ACK lysis (0.15 M NH_4_Cl, 1 mM KHCO_3_, and 0.1mM Na_2_EDTA) of whole blood. Acquisition was performed on a CytoFlex LX (Beckman Coulter) and analysis via FlowJo V10 (Becton Dickinson).

### RNA-Sequencing

The RNA-Seq libraries were generated from purified MkPs from young or old FlkSwitch mice. RNA-Seq libraries were generated using Nextera Library Prep, as we have previously done (Byrne et al. 2017; Beaudin et al. 2016; Poscablo et al. 2021). Libraries were validated using the Bioanalyzer (Agilent 2100), sequenced using Illumina HiSeq 4000 as Paired-end reads at the QB3-Berkeley Genomics at University of California Berkeley and DESeq analysis was done with the help of Dr. Sol Katzman at the UCSC Bioinformatics Core.

### Clustering and Analysis of RNA-Seq data

Normalized RNA abundance counts where extracted from DESeq performed as described above and used to calculate Principal Components (PCs) with the ‘prcomp()’ function in ‘R’ with ‘rank’ set to 50, resulting in 50 components that summate to 100% of the observed variance in the data. PCs 1 and 2 were plotted and used to calculate 1) centroids for each cell type by averaging the sample values in PC1 and PC2 space and 2) Euclidean distances from each centroid to the one calculated for Old HSCs. Furthermore, these normalized counts were scaled to Z-scores using the ‘scale()’ function in ‘R’ with ‘center’ set to ‘TRUÈ. Z-scores were used to create a K-means clustered heatmap with ‘ComplexHeatmap’ in ‘R’. Heatmaps were created on subsets of genes as described in relevant figures with log2FoldChange or Principal Component loading displayed as bar plots where relevant.

### Laser-induced cremaster arteriole thrombosis model

The laser-induced thrombosis assays were performed as previously described (Tourdot & Holinstat 2017; Yeung et al. 2016; Reheman et al. 2009; Adili et al. 2017). Briefly, the cremaster muscle of anesthetized Young (8-12 weeks of age) or Old (>22 months of age) FlkSwitch mice were prepared under a dissecting microscope with constant superfusion of 37°C bicarbonate-buffered saline. Injury of the cremaster arterioles (30-50 um diameter) was performed by a laser ablation system (Ablate! photo-ablation system; Intelligent Imaging Innovations, Denver,CO). Multiple laser injuries were performed in each mouse, with each new injury induced upstream of prior injuries. Images of thrombus formation were taken at 0.2-s intervals using a Zeiss Axio Examiner Z1 fluorescent microscope with a 6x3 objective and a high-speed sCMOS camera. The two populations of Plts were distinguished by the expression of Tomato or GFP fluorescence. Images were analyzed using Slidebook 6.0 (Intelligent Imaging Innovations). Alexa Fluor 647-conjugated anti-fibrin (0.3 μg/g) administered by a jugular vein cannula prior to vascular injury. All captured images were analyzed for the change of fluorescent intensity over the course of thrombus formation after subtracting fluorescent background defined on an uninjured section of the vessel using the Slidebook program.

### Platelet-Leukocyte Aggregation Analysis

10 μl of heparinized blood was aliquoted into an antibody cocktail containing CD41-APC, B220-BV605, Ter119-A700, GR-1 PB and incubated at room temperature in the dark for 20 minutes. In samples to be stimulated by Thrombin, Thrombin (0.1Units/ml) was added to the cocktail and incubated for 5 minutes at room temperature and protected from light. 500 μl 1-step Fix/Lyse Solution (Invitrogen # 00-5333-57) was added to all samples and allowed to incubate another 30 minutes in the dark at room temperature. Samples were analyzed by flow cytometry within 2 hours of obtaining blood samples.

### Platelet Activation and Glycoprotein Expression Analysis

Platelets were obtained from whole blood collected in EDTA coated capillary tubes, washed with Tyrode’s buffer, and examined under resting conditions or after activation with 1 U/µl Thrombin (Sigma) or with 10 µM adenosine diphosphate (ADP, Sigma) for 5 minutes at 37°C, followed by 5 minutes at room temperature. Platelets were stained with anti-CD9, anti- αIIb/β3 (JON/A, Emfret), and anti-P-Selectin. Glycoprotein expression on activated platelets were measured by flow cytometry.

### Platelet Depletion and Cell Cycle Analysis

Acute thrombocytopenia was induced in Y and O FlkSwitch mice by depleting Plts with a single injection of anti-GPIbα (2 µg/g) intraperitoneally (R300, Emfret). Retro-orbital (RO) bleeds using heparin-coated capillary tubes were performed daily to monitor Plt recovery. Plts were measured by flow cytometry and defined as single, FSC^low^Ter119-CD61+ Tom/GFP+ cells. In a separate cohort of mice, HSPCs in the BM was analyzed 24 hours post anti-GPIbα treatment. In the cell cycle experiments, EdU (50mg/kg) was administered intraperitoneally and concurrent with anti-GPIbα administration. Following a 24-hour EdU pulse, HSCs, MPPs, and MkPs were purified by flow cytometry and EdU incorporation analyses were performed according to the manufacturer’s protocols (Click-iT® EdU Flow Cytometry Assay Kits, Life Technologies).

### Quantification and Statistical Analysis

Number of experiments, n, and what n represents can be found in the legend for each figure. Statistical significance was determined by two-tailed unpaired T-test, unless noted in Figure Legends. All data are shown as mean ± standard error of the mean (SEM) representing at least three independent experiments.

### Data availability

The datasets generated in the current study are available in the Gene Expression Omnibus (GEO), accession number GSE166704. https://www.ncbi.nlm.nih.gov/geo/query/acc.cgi?acc=GSE166704

