## Supplementary material for "An age-specific platelet differentiation path from hematopoietic stem cells contributes to exacerbated thrombosis": Poscablo Supplemental Data

**Figure S1. Gating strategies for hematopoietic stem and progenitor populations in young and old FlkSwitch bone marrow.**

**A.** Representative FACS plots of HSC, MPP, GMP, CMP, and MEP.

**B.** Representative FACS plots of Common Lymphoid Progenitors (CLP).

**C.** Representative FACS plots of MkPs, CFU-E, pCFU-E, pGMP, pMegE, and “GMP”.

**D.** Increased HSC and MkP frequencies in old mice. Frequency of HSCs and MkPs in Lineage- BM cells. Data represent 6 independent experiments, n = 8 young mice, n = 18 old mice. Statistics: t-test. ** P< 0.005, *** P< 0.0005

**E.** FACS plots representing Tom and GFP expression by HSCs and classical myeloid progenitors, and CLPs from panel A and B, respectively.

**F.** FACS plots representing Tom and GFP expression by MkPs and alternatively defined erythromyeloid progenitors from panel C.

**Figure S2. Gating strategies for tissue-resident macrophages in FlkSwitch mice across tissues.**

**A.** FACS plots representing the GFP labeling in microglia in the brain of young and old FlkSwitch mice: CD45+F4/80$^{hi}$CD11b$^{hi}$Ly6g−CD11c−.

**B.** FACS plots representing the GFP labeling in tissue resident alveolar macrophages in the lungs of young and old FlkSwitch mice: CD45+F4/80$^{hi}$CD11b$^{mid}$ SiglecF+.

**Figure S3. Gating strategies for Multipotent Progenitor subfractions in the FlkSwitch mice during aging.**

**A.** FACS plots representing the GFP and Tom expression by subfractions of Multipotent Progenitors: MPP2 (Lin-cKit+Sca1+Flk2-CD48+SLAM+), MPP3 (Lin-cKit+Sca1+Flk2-CD48+SLAM-), and MPP4 (Lin-cKit+Sca1+Flk2+CD48+SLAM-).

**B.** The frequency of MPP subpopulations do not change substantially upon aging. Data represents 3 independent experiments, n = 5 young mice, n = 5 old mice. Statistics: t-test. Comparisons were not statistically different. Y, young; O, old.

**Figure S4. Old HSCs poorly reconstitute nucleated cells and platelets compared to young HSCs upon transplantation.**

Percentages of donor chimerism in young (A) or old (B) recipient mice transplanted with oHSCs from FlkSwitch mice demonstrate feeble long-term reconstitution of nucleated cells (GM, B, and T cells) and Plts compared to yHSCs. Data represent mean ± SEM of 5 independent experiments with n= 15 Y-Y mice, n= 21 O-Y mice, n= 13 Y-O mice, n= 13 O-O mice. Statistics: unpaired two-tailed t-test between total cell (black) or Plt (magenta) chimerism by oHSCs and yHSCs. *p<0.05, **p<0.005, ***p<0.005.

**Figure S5. Heatmaps of RNAseq analysis.**

**A.** Kmeans-clustered heatmap of gene expression Z-score for top 30 genes by absolute loading in Principal Component 1 (top) or 2 (bottom) derived for all samples.

**B.** Kmeans-clustered heatmap of gene expression Z-score for genes with MkP- or HSC-specific expression according to the GEXC database. MkP- or HSC- specific genesets were targeted from GEXC with the search term of “active in MkP, while inactive in all other cells" and “active in HSC, while inactive in all other cells", respectively.

**Figure S6. Tom+ MkPs from old FlkSwitch mice have greater myeloid reconstitution potential compared to GFP+ Young or Old MkPs.**

**A.** Analysis of donor-derived erythroid cells, GM, B and T cells in the peripheral blood of recipients presented as percent donor chimerism. Tom+ oMkPs demonstrated greater contribution to erythroid (beyond day 30 post-transplant) and GM (all post-transplant timepoints) donor-to-host chimerism in the recipient mice compared to both GFP+ yMkPs and GFP+ oMkPs. Little to no B and T cell chimerism was observed. Data are from the same cohorts of mice presented in Figure 4, and represent mean ± SEM of 3 independent experiments, n = 6 GFP+ yMkP recipients, n = 4 GFP+ oMkP recipients, and n = 13 Tom+ oMkP recipients. Statistics: unpaired two-tailed t-test. T-tests between GFP+ yMkP and GFP+ oMkP were not statistically significant. Erythroid chimerism: T-tests between GFP+ yMkP and Tom+ oMkP #p<0.05 at Day 14 and Day 28. T-tests between GFP+ oMkP and Tom+ oMkP *p<0.05, **p<0.005, ***p<0.0005.

**B.** Phenotypic MkPs can be stratified based on CD48 expression. Representative flow cytometry plot (left) and quantification of MkP CD48 subtypes (right). Data represent more than 20 independent experiments, n=39 young WT mice. Statistics: unpaired two-tailed t-test, ****P<0.0001.

**C.** Equivalent in vitro growth capacity by CD48+ or $CD48^{lo/-}$ MkPs. Quantification of cell expansion revealed no difference between young CD48+ or $CD48^{lo/-}$ MkPs. Data represent 7 independent experiments, n=13 CD48+ yMkPs and n=8 $CD48^{lo/-}$ yMkPs (each data point is the average of three replicate wells). Statistics: unpaired two-tailed t-test, ns = not statistically significant.

**D.** Equivalent Plt reconstitution by CD48+ or $CD48^{lo/-}$ yMkPs. Data represent mean ± SEM of 5 independent experiments, n=8 CD48+ yMkPs, n=3 $CD48^{lo/-}$ yMkPs, and n=8 Sham. Statistics: Repeated measures two-way ANOVA with the Geisser-Greenhouse correction, adjusted for multiple comparisons by Tukey's ad hoc test.

**E.** Similar to GFP+ yMkPs, CD48+ or $CD48^{lo/-}$ yMkPs from wt mice contribute minimally to GM, B, and T cell reconstitution following transplantation. $CD48^{lo/-}$ yMkPs specifically contribute to transient erythroid cell output. Percent donor chimerism was assessed via sequential tail bleeds at the indicated time points. Data from same cohort as **D.** **P<0.01, ****P<0.0001.

**Figure S7. The age-specific Tom+ Plts contribute to exaggerated clot formation in old FlkSwitch mice.**

Images of uninjured FlkSwitch mice (A) and laser-induced clot formation (B) in Young and Old FlkSwitch mice. Thrombi in Old mice contain both Tom+ and GFP+ cells, whereas thrombi in young mice are exclusively GFP+.

**Figure S8. Gating strategy for Platelet-Leukocyte Aggregation Analysis of Young and Old Mice.** Gating strategy of platelet-leukocyte aggregates in vitro analysis to determine frequency of platelet aggregates with distinct leukocyte subtypes within stimulated murine blood. Gr1+CD41+ events (A) and B220+CD41+ events (B). Stimulation by thrombin (0.1 U/mL) increased the double-positive events in young and old blood. Platelet-leukocyte aggregation remained higher in old blood compared to young blood, with and without stimulus by thrombin.

.

**Movie S1.**

In vivo laser ablation injury and real-time monitoring of thrombus formation in a Young FlkSwitch mouse. Note formation of a small clot consisting of GFP+, but not Tom+, cells.

**Movie S2.**

In vivo laser ablation injury and real-time monitoring of thrombus formation in an Old FlkSwitch mouse. Note formation of a large clot consisting of both Tom+ and GFP+ cells.

### Supplemental Figure 1

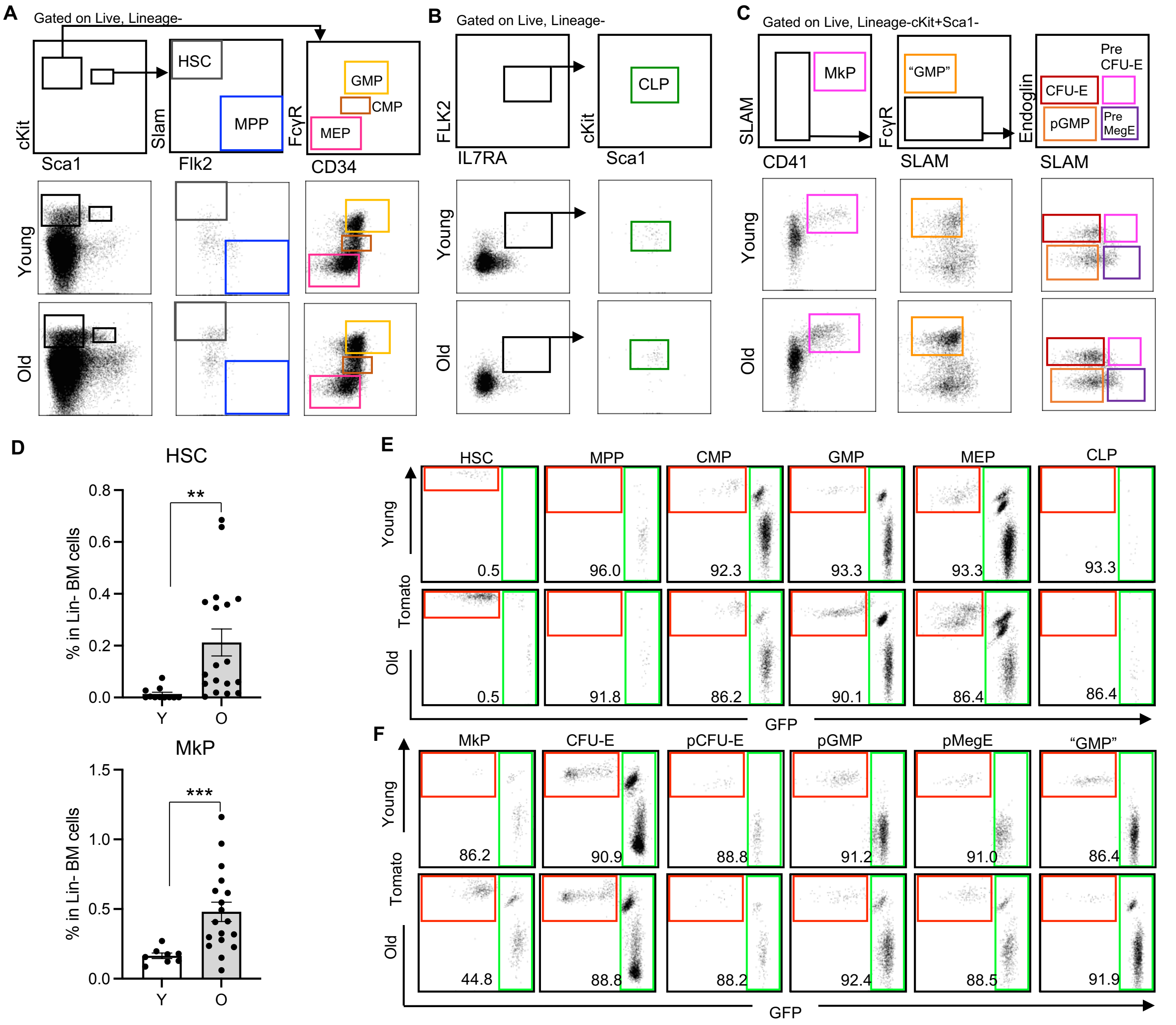

### Supplemental Figure 2

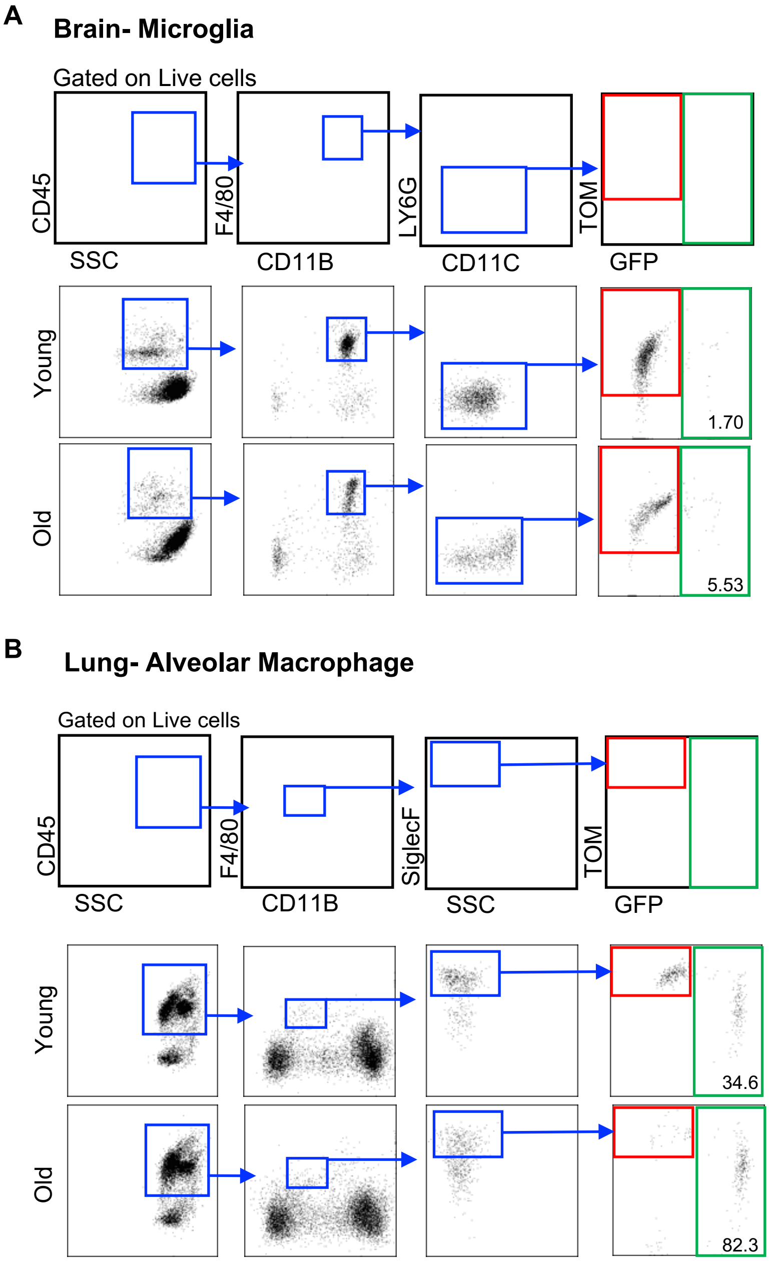

### Supplemental Figure 3

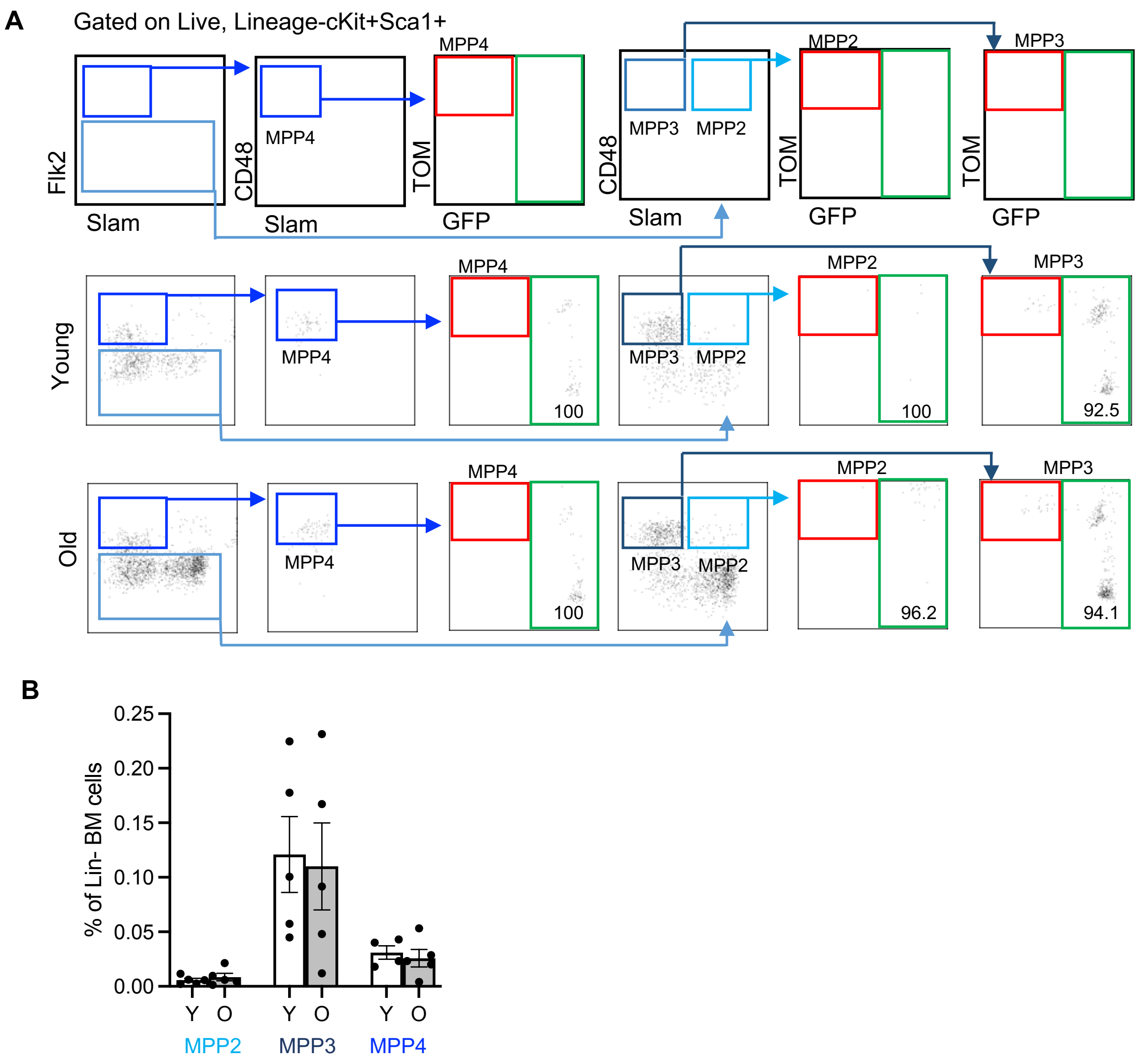

### Supplemental Figure 4

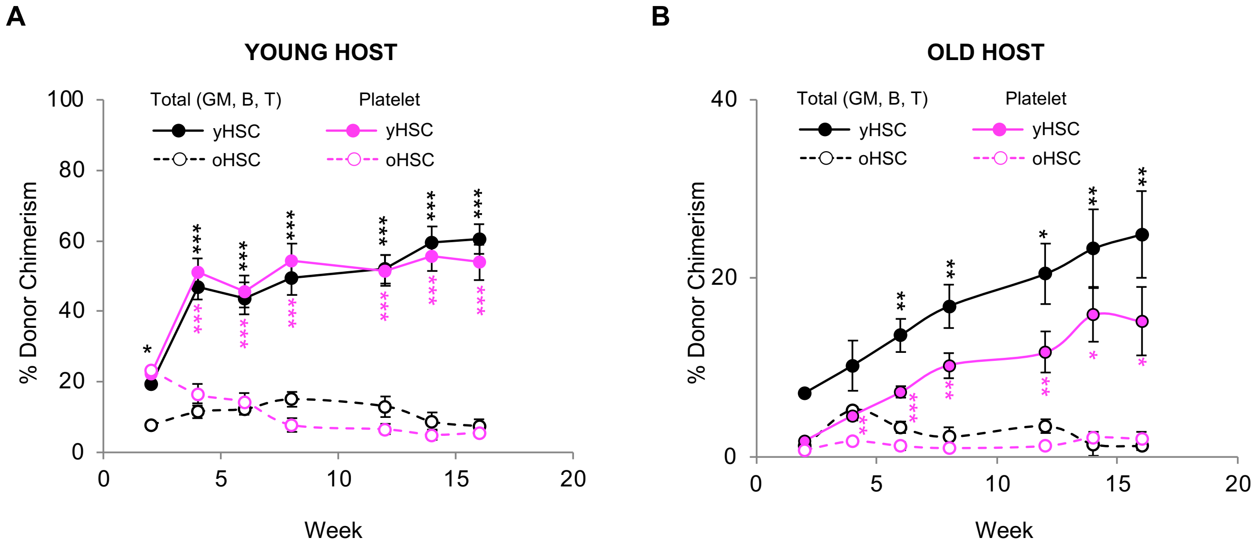

### Supplemental Figure 5

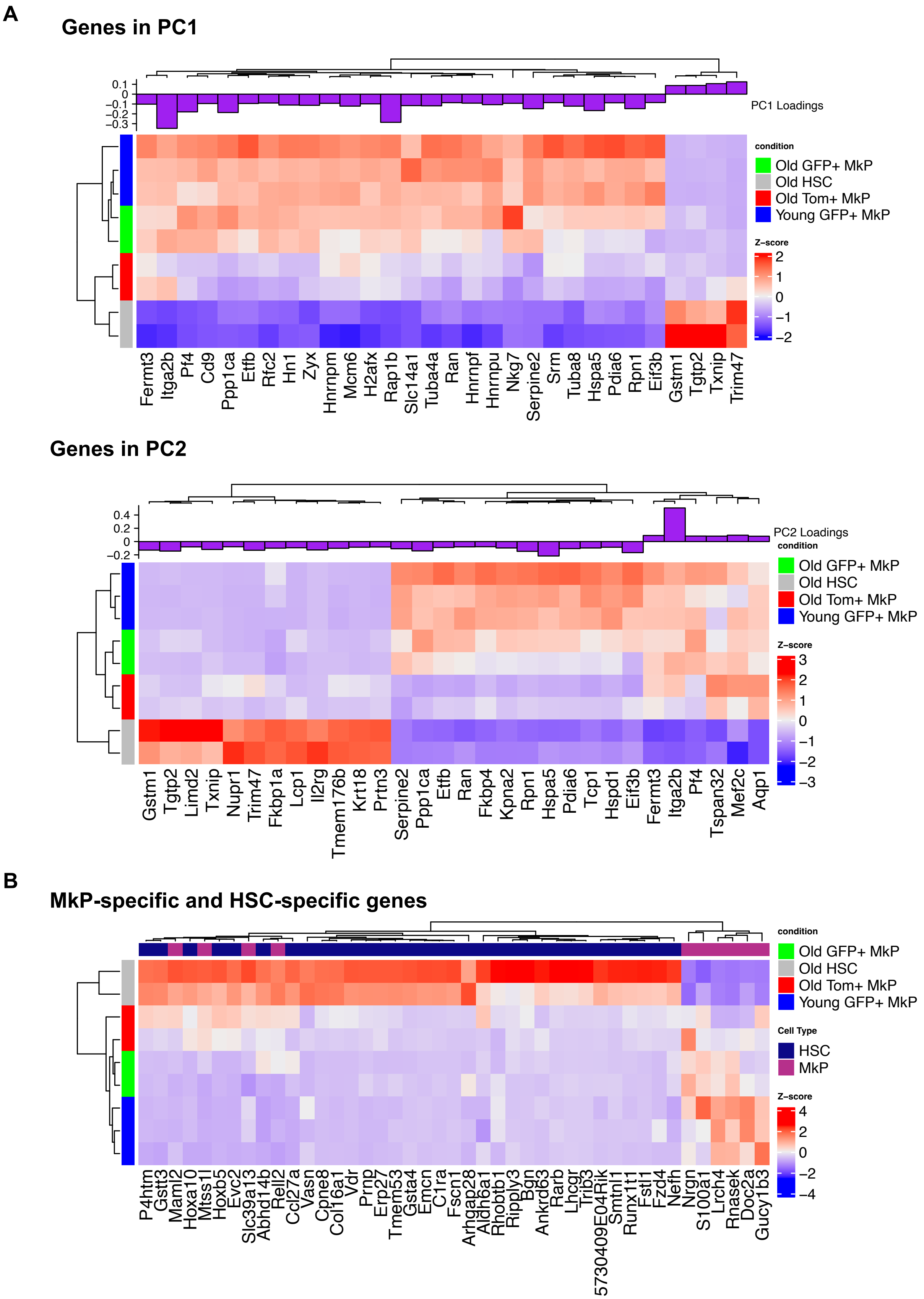

### Supplemental Figure 6

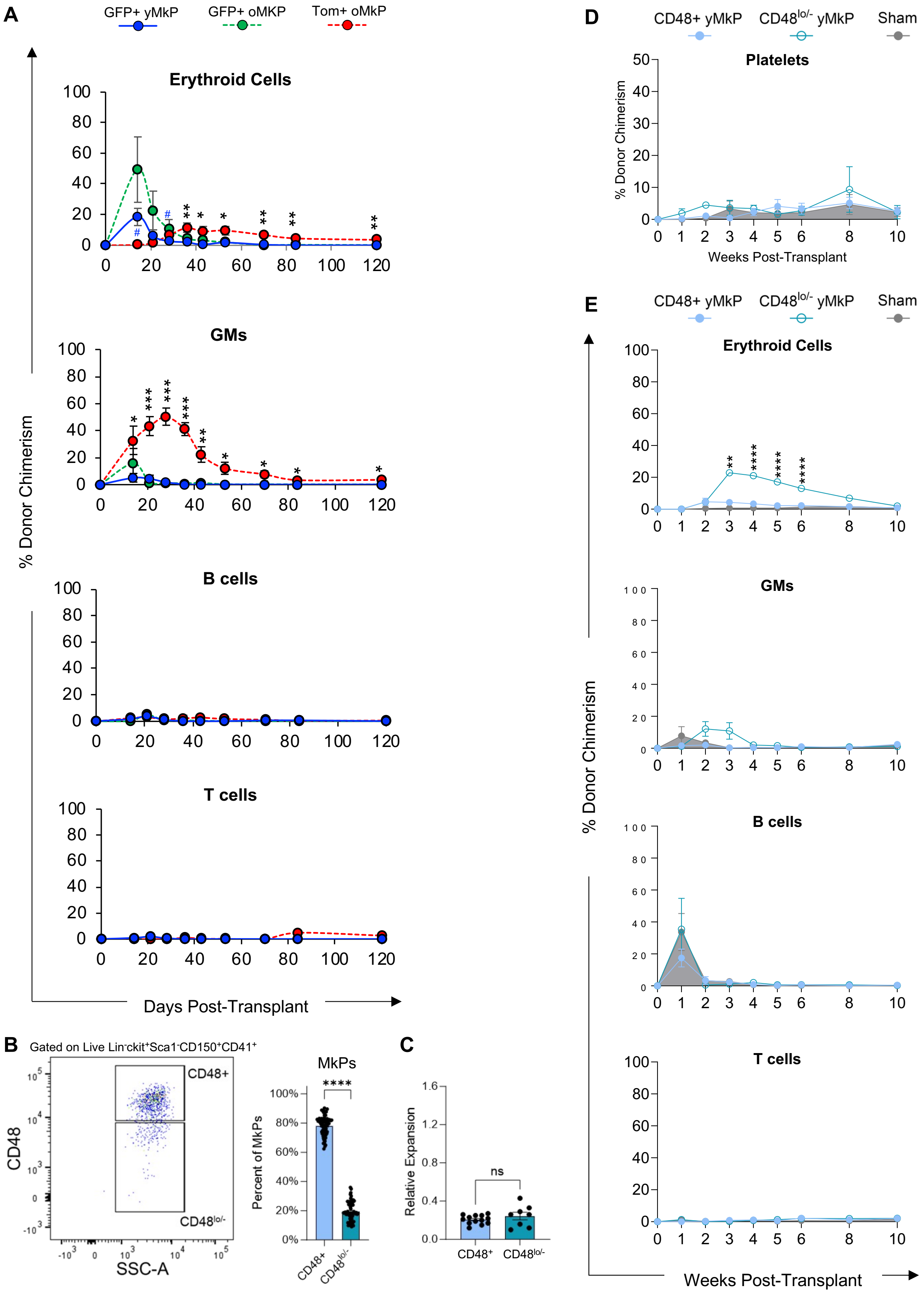

Supplemental Figure 7

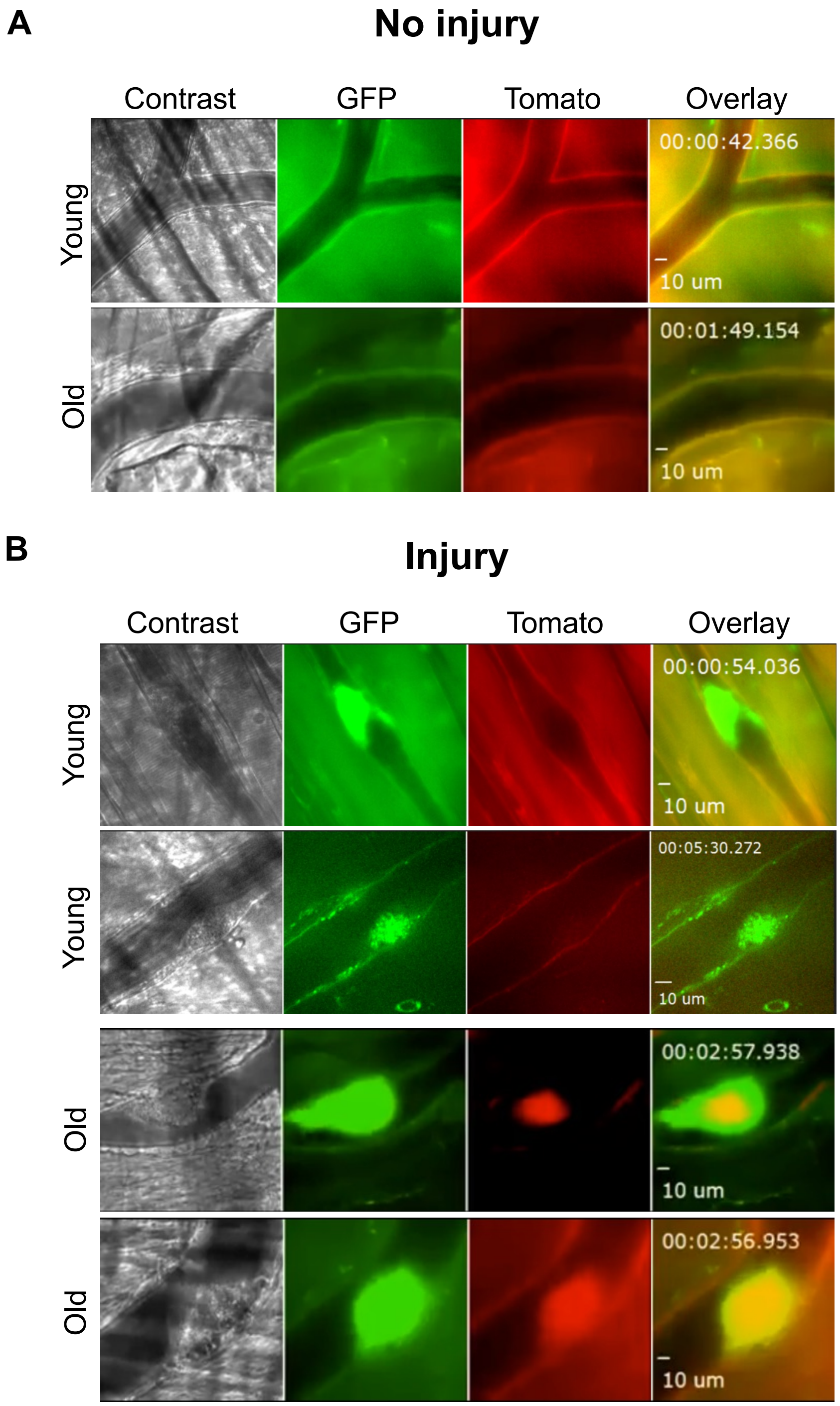

#### Supplemental Figure 8

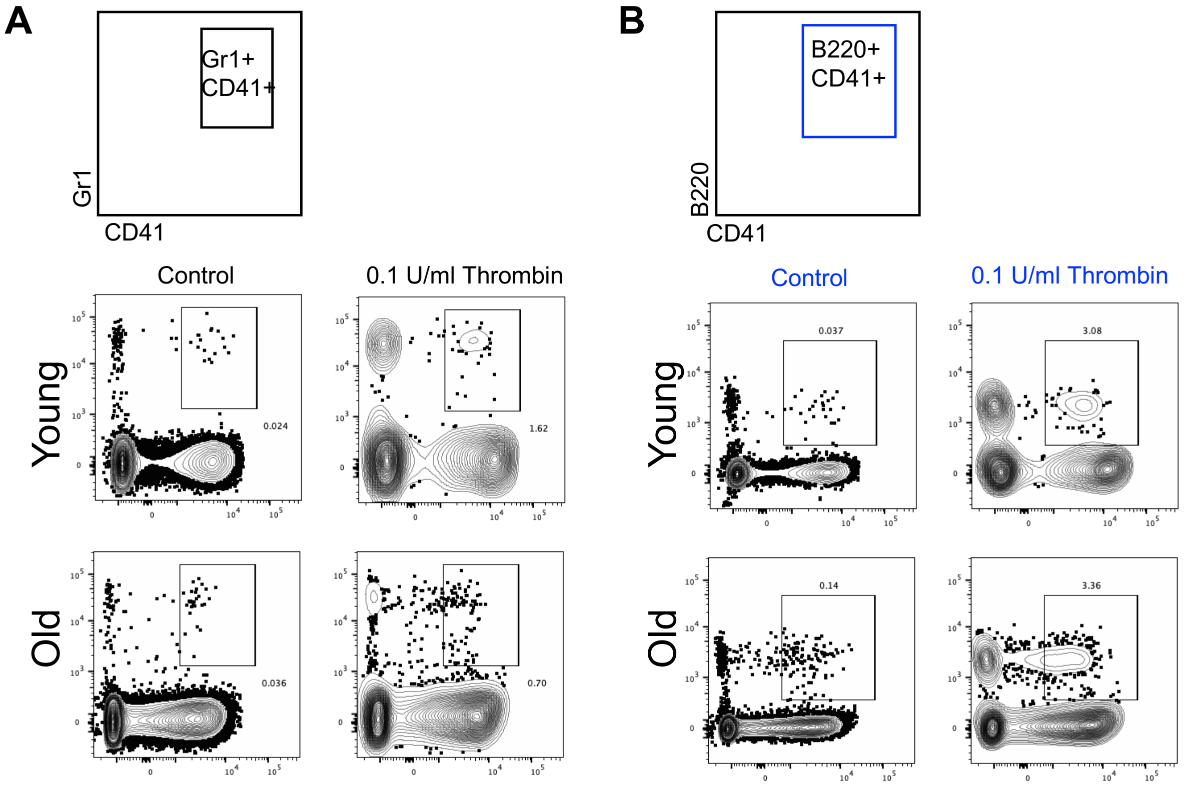

#### Supplemental Movie 1

mp4 files: https://drive.google.com/drive/folders/1Ix50j4AVRqp0axve8m7kr8N1WTGfXMpy

##### Young 1_example in Fig.5

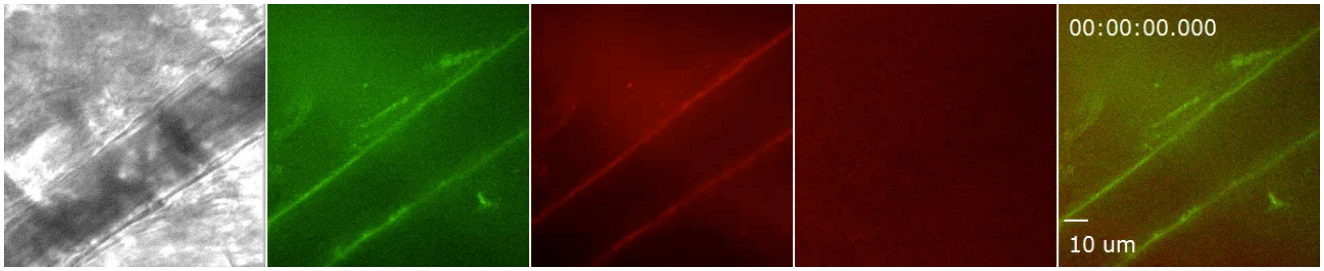


##### Young 2

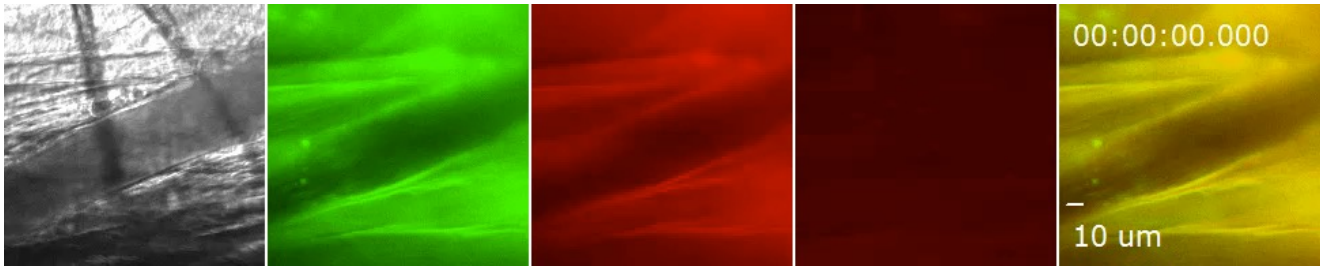


##### Young 3

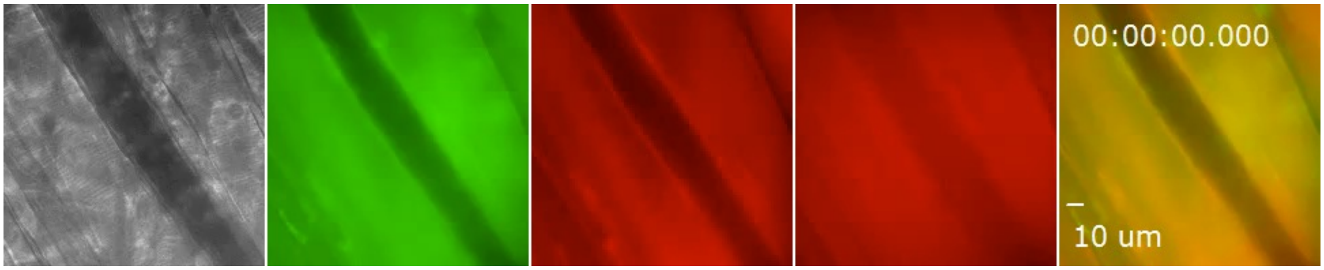

Supplemental Movie 2
mp4 files: https://drive.google.com/drive/folders/1Ix50j4AVRqp0axve8m7kr8N1WTGfXMpy

Old 1_example in Fig.5

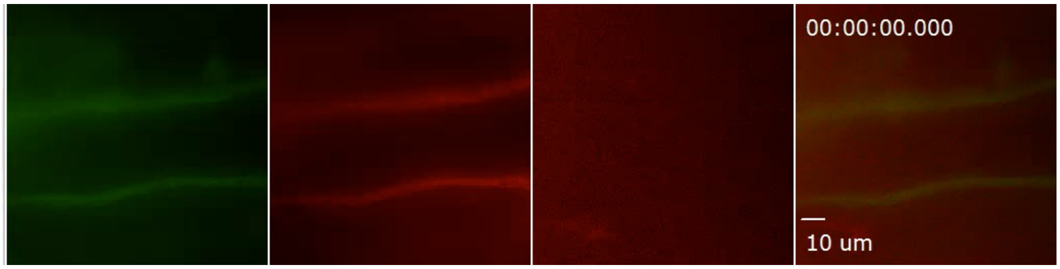


Old 2

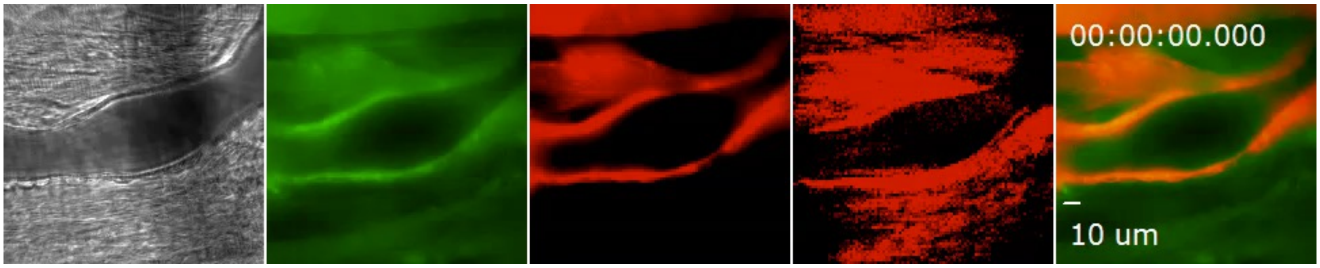


Old 3

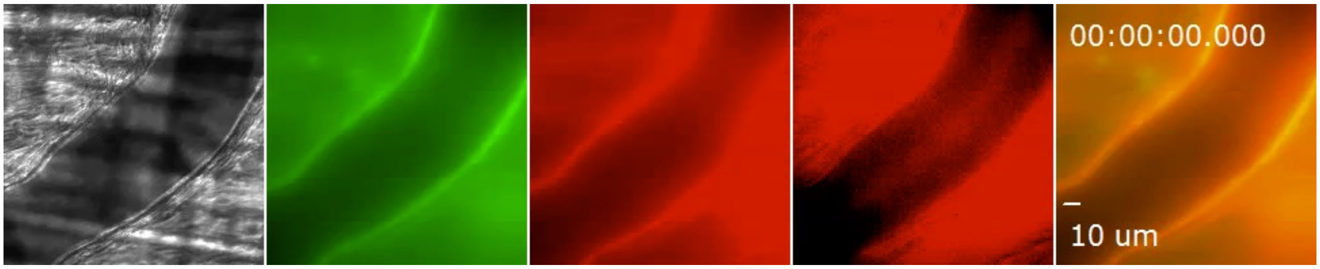
